# SWAP1-SFPS-RRC1 splicing factor complex modulates pre-mRNA splicing to promote photomorphogenesis in Arabidopsis

**DOI:** 10.1101/2022.04.26.489584

**Authors:** Praveen Kumar Kathare, Ruijiao Xin, Abirama Sundari Ganesan, Viviana M. June, Anireddy S. N. Reddy, Enamul Huq

## Abstract

Light signals perceived by a group of photoreceptors have profound effects on the physiology, growth, and development of plants. The red/far-red light absorbing phytochromes modulate these aspects by intricately regulating gene expression at multiple levels. Previously, we reported that two splicing factors SFPS (SPLICING FACTOR FOR PHYTOCHROME SIGNALING) and RRC1 (REDUCED RED LIGHT RESPONSES IN CRY1CRY2 BACKGROUND 1), interact with photoactivated phyB to regulate light-mediated pre-mRNA alternative splicing (AS). Here, we report the identification and functional characterization of an RNA binding splicing factor, SWAP1 (SUPPRESSOR-OF-WHITE-APRICOT/SURP RNA-BINDING DOMAIN-CONTAINING PROTEIN1). Loss-of-function *swap1-1* mutant is hyposensitive to red light and exhibits a day light-independent early flowering phenotype. SWAP1 physically interacts with both SFPS and RRC1 in a light-independent manner and forms a ternary complex. In addition, SWAP1 also physically interacts with photoactivated phyB and colocalizes with nuclear phyB photobodies. Deep RNA-seq analyses show that SWAP1 regulates the gene expression and pre-mRNA alternative splicing of a large number of genes including those involved in plant responses to light signaling. A comparison with SFPS- and RRC1-regulated events shows that all three splicing factors coordinately regulate the alternative splicing of a subset of genes. Collectively, our study uncovered the function of a new splicing factor, which interacts with photoactivated phyB, in modulating light-regulated development in plants.

**SIGNIFICANCE:** Regulation of transcription and pre-mRNA alternative splicing is essential for the transcript diversity and modulation of light signaling in plants. Although several transcription factors involved in light signaling have been discovered and characterized in-depth, only a few splicing factors have been shown to be involved in the regulation of light signaling pathways. In this study, we describe the identification and characterization of a new splicing factor SWAP1, which interact with two previously characterized splicing factors, SFPS and RRC1, forming a ternary complex. We show that, like SFPS and RRC1, SWAP1 also interacts with photoactivated phyB, and consistently, *swap1* seedlings are hyposensitive to red light. SWAP1 modulates alternative splicing of a large number of genes and a subset of these genes are coordinately regulated by SFPS, RRC1 and SWAP1. These results highlight the importance of not only the transcription factors but also the phyB-interacting splicing factors in light-regulated plant development.

## INTRODUCTION

Light functions not only as an energy source for photosynthesis but also as an environmental signal, which modulates the growth and development throughout the plant life cycle. To perceive and respond to surrounding light, plants contain an array of photoreceptors with distinct and/or overlapping wavelength perception. These photoreceptors collectively sense the light intensity, color and duration to optimize growth and development of plants (1, 2). One such photoreceptor, ubiquitous across the plant and bacterial kingdom is ‘phytochrome’ (phy), which perceives and responds to red/far-red wavelength of the light spectrum (3, 4). In Arabidopsis, phys are encoded by a multigene family consisting of 5 different genes (*PHYA to PHYE*) (5). In dark-grown plants, phys are predominantly localized to the cytosol in the inactive Pr form, and upon red light illumination, they are photoconverted to biologically active Pfr form and translocate into the nucleus to form discrete nuclear bodies called photobodies (PBs) (3, 4). These PBs are membraneless subnuclear dynamic structures, whose precise molecular functions are yet to be understood. Nevertheless, based on several molecular, biochemical, and imaging studies, it has been hypothesized that these PBs are probably molecular structures within which phys interact with diverse target proteins to initiate appropriate signaling cascades (6, 7). One group of proteins, which has been recently identified to be interacting with photoactivated phyB and colocalized to PBs are splicing factors, that regulate light-induced pre-mRNA alternative splicing (AS) (8).

Pre-mRNA AS is an essential process in eukaryotes, which increases the complexity of gene expression by generating multiple forms of mature mRNAs from a single multi-exon/intron pre-mRNA (4, 9–11). Both internal and external factors modulate the pre-mRNA AS to optimize the mRNA diversity (12–17). A recent comprehensive analysis of AS has revealed that about 79% of multi-exonic protein-coding pre-mRNAs undergo AS in response to different stimuli, suggesting a high prevalence of AS phenomenon in Arabidopsis (18). AS is a tightly regulated and multistep biochemical process accomplished by a highly conserved, dynamic, and flexible macromolecular ribonucleoprotein complex called spliceosome (11). The functional spliceosome complex consists of a core and a group of *trans*-acting factors (also known as auxiliary splicing regulatory proteins). The core is made up of approximately 200 proteins plus a set of five small nuclear ribonucleoproteins (snRNPs; U1, U2, U4, U5 and U6), while *trans*-acting factors such as SR (Serine/Arginine-rich) proteins and hnRNPs (heterogeneous nuclear ribonucleoproteins) variably interact with the core to guide spliceosome complex to modulate appropriate AS event(s) (10, 19, 20). Pre-mRNAs destined for AS contain a set of *cis*-acting elements, which include exonic and intronic splicing enhancers (ESEs and ISEs) as well as exonic and intronic splicing silencers (ESSs and ISSs) (11, 21). Interactions between *cis*-acting elements and *trans*-acting factors eventually define the AS event(s) to generate the mature mRNA. Positively acting *trans*-acting factors such as SR proteins typically bind to ESEs/ISEs promoting splicing, while negatively acting *trans*-acting factors such as hnRNPs typically bind to ESSs/ISSs suppressing pre-mRNA splicing. Thus, a cascade of protein-protein and protein-RNA interactions in response to a range of stimuli modulate the optimal pre-mRNA splicing (10, 11, 19, 21, 22).

The critical role of phys in the modulation of global gene expression pattern is well documented and in the past few years, it has been shown that the photoactivated phys also regulate pre-mRNA AS (8, 9, 22–27). Amongst the most crucial AS targets of photoactivated phys are pre-mRNAs encoding several SR proteins and U1/U2 snRNPs, which in turn further control the AS of many target pre-mRNAs (17, 23). Recently, we have shown that at least two *bona fide* splicing factors, SFPS (SPLICING FACTOR FOR PHYTOCHROME SIGNALING) and RRC1 (REDUCED RED-LIGHT RESPONSES IN CRY1CRY2 BACKGROUND1) interact with photoactivated phyB and modulate pre-mRNA splicing of a large number of genes to optimize photomorphogenesis in Arabidopsis (28, 29). In this study, we describe the identification and characterization of a novel SFPS interacting protein partner, SWAP1 (Suppressor-of-White-APricot1; At4g31200), a putative RNA-binding protein, which forms a ternary complex with SFPS and RRC1 to modulate plant responses to red light by regulating AS of pre-mRNAs of a subset of light-regulated genes.

## RESULTS

### *swap1-1, like sfps-2 and rrc1-3*, is hyposensitive to red light

Recently we have shown that SFPS and RRC1 are involved in light-regulated pre-mRNA AS and differential gene expression to modulate photomorphogenesis in Arabidopsis (28, 29). SFPS was first identified in a forward genetic screen of mutants defective in red light signaling and subsequently, RRC1 was identified as one of the interacting partners of SFPS in an IP-MS (Immunoprecipitation followed by mass spectrometry) analysis. The same IP-MS study also identified SWAP1 as another potential interacting partner of SFPS (Fig. S1A and B). SWAP1 is a ~73kda protein and an *in-silico* data analysis categorized SWAP1 as an RNA binding protein containing SWAP and RPR (regulation of nuclear pre-mRNA) domains in its N- and C-termini, respectively (Fig. S1C).

To understand the biological function of SWAP1, we obtained a homozygous T-DNA insertional mutant (*swap1-1*; SALK_027493; Fig. S2A and B) from ABRC and examined seedling phenotypes under dark, continuous red, far-red, and blue light conditions. *swap1-1* displayed wild-type sensitivity under dark, far-red, and blue light conditions, while it exhibited long hypocotyl phenotypes under red light (Fig. 1A and B, Fig. S2C and D). In addition, *SWAP1_pro_:SWAP1-GFP/swap1-1* transgenic lines grown under similar red light conditions complemented the *swap1-1* mutant phenotype (Fig. S3A-D). Although SWAP1 regulates red light responses, the abundance of *SWAP1* mRNA and protein was not altered in response to prolonged red light illumination (Fig. S3E and F).

**Figure 1:**
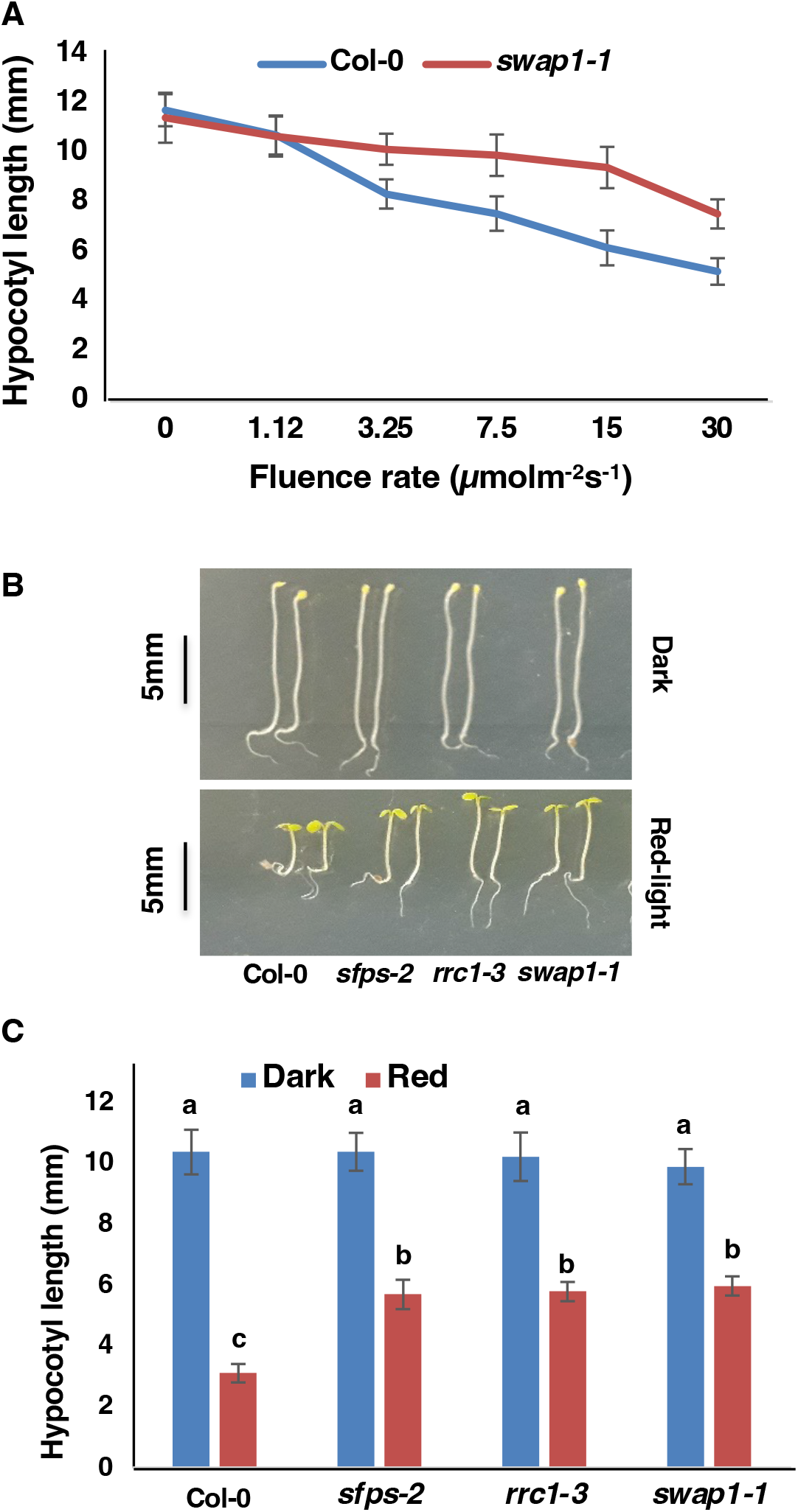
*swap1-1* mutant is hyposensitive to red light. (A) Quantification of the hypocotyl length of 4-day-old seedlings grown under either continuous dark or continuous red light of different fluence rate. (B and C) Digital images of representative seedlings (B) and bar graph showing hypocotyl lengths (C) of different genotypes grown under either continuous dark or continuous red light (7μmolm^-2^s^-1^) for 4 days. Bars indicate mean length in mm and error bars indicate standard deviation. Statistical significance among different genotypes was determined using single factor ANOVA followed by Tukey’s post hoc analysis and is indicated by different letters.

To test whether *SWAP1* participates in the modulation of vegetative to reproductive growth in Arabidopsis, the flowering time of wild-type and *swap1-1* mutant was quantified and compared by counting the number of rosette leaves and days-to-flower under both short-day (SD; 8hrs light: 16hrs dark) and long-day (LD; 16hrs light: 8hrs dark) conditions. The *swap1-1* mutant plants flowered earlier than wild-type plants, both under SD and LD conditions. On the other hand, *SWAP1_pro_:SWAP1-GFP/swap1-1* transgenic plants flowered at the same time as wild-type plants (Fig. S4 and S5). Taken together, these data suggest that SWAP1 is one of the essential regulatory components that modulates red light signaling and day-light-independent flowering in Arabidopsis.

### SWAP1 interacts with SFPS and RRC1 and forms a ternary complex

As SWAP1 was found to co-immunoprecipitate with SFPS, we investigated whether SWAP1 is a bona fide interacting partner of SFPS and RRC1 through a series of *in vitro* and *in vivo* protein-protein interaction studies. Yeast two-hybrid and *in vitro* pull-down assays using full-length SWAP1, SFPS and RRC1 indicated that the SWAP1 physically interacts with both SFPS and RRC1 (Fig. S6). *In vivo* Co-IP assay performed using the dark and red light-treated samples indicated that the SWAP1 interacts with both SFPS and RRC1 in a light-independent manner (Fig. 2A and B). Furthermore, Co-IP assays also revealed that the presence or absence of SFPS or RRC1 does not alter the strength of interaction between SWAP1 and RRC1 or SWAP1 and SFPS, respectively (Fig. 2A and B). Since each of the three proteins physically interact with each other, these three proteins may form a functional ternary complex to modulate pre-mRNA splicing. To test the formation of the ternary complex, *in vitro* pull-down assay was performed using MBP-RRC1 as a bait protein and GST-SFPS and an increasing concentration of GST-SWAP1 as prey proteins. The results show that MBP-RRC1 interacts with both GST-SFPS and GST-SWAP1 simultaneously forming a ternary complex (Fig. 2C).

**Figure 2:**
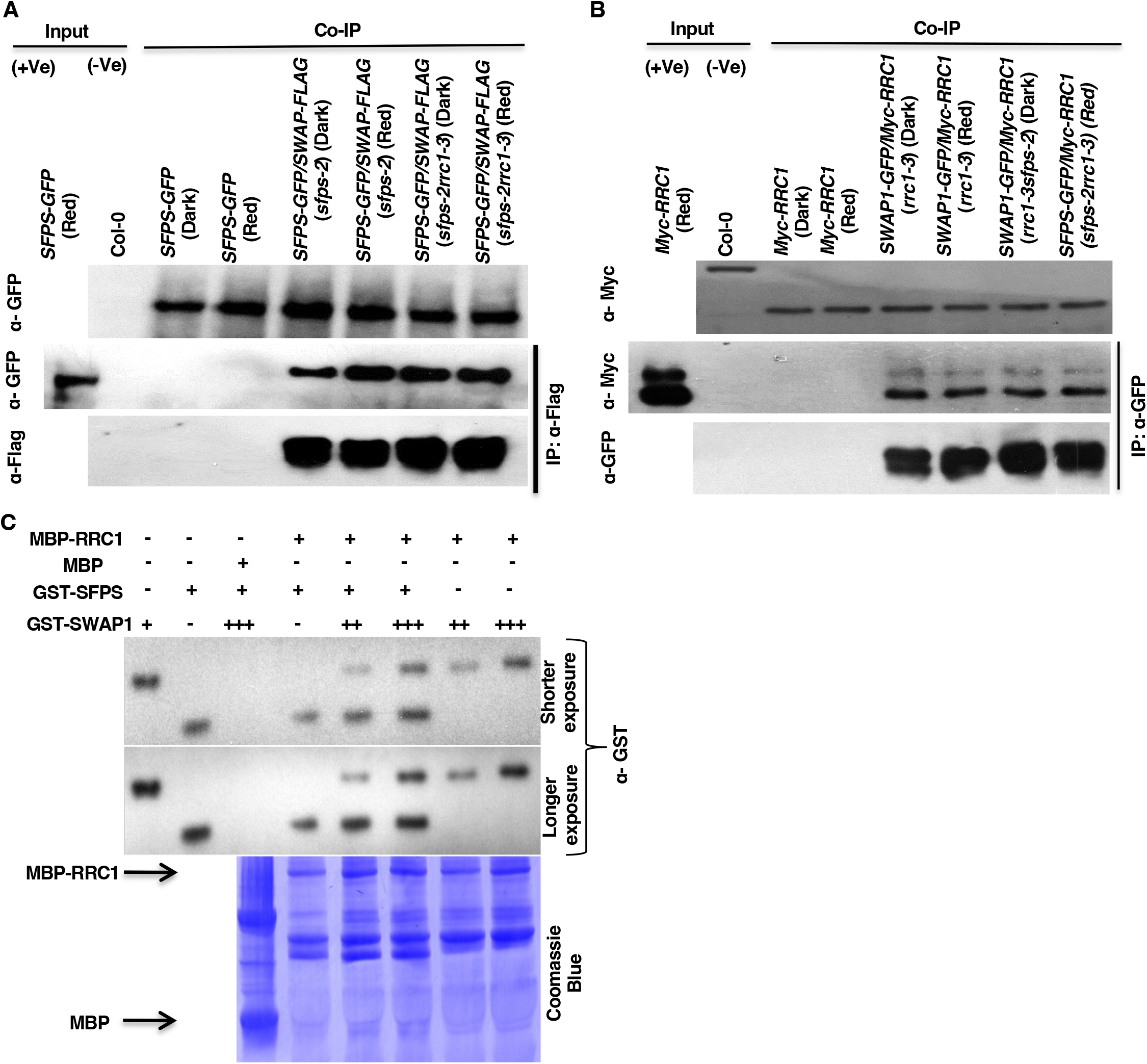
SWAP1 interacts with SFPS and RRC1 and forms a ternary complex. (A and B) SWAP1 interacts with SFPS (A) and RRC1 (B) *in vivo*. Four-day-old seedlings were either kept in the dark or exposed to the constant red light for 6 hrs (7μmolm^-2^s^-1^) and total protein was extracted in native extraction buffer. 1μg of appropriate antibody bound to Dynabeads (protein A) was added to the protein mixture and incubated for 3hrs at 4°C. After the incubation, beads were pelleted and washed multiple times and proteins were separated on 8-10% SDS-PAGE. The presence or absence of specific proteins was detected following western blotting with appropriate antibodies. (C) SWAP1, SFPS and RRC1 proteins form a ternary complex *in vitro*. Bacterially expressed and amylose resin-bound MBP and MBP-RRC1 were used as bait proteins to pull-down GST-SFPS and GST-SWAP1 prey proteins. For GST-SFPS and GST-SWAP1, += ~1μg, ++= ~2μg and +++= ~3μg of protein. The pull-down reaction was incubated for 3hrs at 4°C on a rotating shaker. After the incubation, beads were washed at least 5 times. Proteins were separated on 10% SDS-PAGE gel, transferred to PVDF membrane and first immunoblotted with α-GST, followed by Coomassie blue staining of the membrane.

Subcellular localization of SWAP1-GFP revealed that SWAP1 localizes exclusively to nucleus and forms discrete nuclear speckles of varying sizes (Fig. S7A). As reported earlier, both SFPS and RRC1 also form discrete nuclear speckles and colocalize with each other (29). Since SWAP1 interacts with both proteins, we examined whether SWAP1 nuclear speckles also colocalize with those of SFPS and RRC1. We prepared SFPS-GFP/SWAP1-mCherry and SWAP1-GFP/RRC1-mCherry double transgenics, and fluorescent confocal imaging was performed using 8-day-old light-grown seedlings. The results show that the SWAP1 nuclear speckles almost always colocalize with nuclear speckles of SFPS and RRC1 (Fig. S7B and C). Taken together, these biochemical and imaging data suggest that SWAP1, SFPS and RRC1 proteins not only colocalize with each other but also physically interact to form a ternary protein complex.

Previously, SFPS and RRC1 were shown to co-localize with 3’ SS (Splice Site) targeting U2-snRNPs associated components such as U2A’, U2AF35A and U2AF65B in the nucleus (28, 29). As SWAP1 interacts and co-localizes with both SFPS and RRC1, we examined whether SWAP1 col-localizes with U2-snRNP-associated components. Double transgenic lines expressing SWAP1-GFP/U2A’-mCherry, SWAP1-GFP/U2AF35A-mCherry and SWAP,1-GFP/U2AF65B-mCherry were prepared and root cells from different developmental regions were observed under confocal fluorescent microscopy. Interestingly, the nuclear speckles of SWAP1-GFP invariably co-localize with the nuclear speckles of U2A’-mCherry, U2AF35A-mCherry and U2AF65B-mCherry (Fig. S8), suggesting that SWAP1 might regulate 3’ SS selection.

### SWAP1 regulates the expression of genes involved in light signaling

Splicing factors are reported to regulate gene expression in response to a specific stimulus (8, 10, 22). As SWAP1 is predicted to be a splicing factor, we examined whether it modulates gene expression by performing deep RNA sequencing of wild-type and *swap1-1* mutant seedlings grown under dark and dark-grown seedlings exposed to 3 hrs of red light. RNA-seq data were first analyzed using the DESeq2 package and then enriched Gene Ontology (GO) terms were determined using the GeneCodis4 online tool. Similar to our earlier observations, a comparison of differentially expressed genes (DEGs) between dark and red light-treated wild type samples identified a total of 6226 DEGs passing the threshold of fold change (FC) of ≥ 1.5 and false discovery rate (FDR) of ≤ 0.05. In contrast, red light irradiated *swap1-1* displayed 7595 DEGs compared to the dark-treated *swap1-1* mutant samples (Fig. S9A, Dataset S1, I and VII). An in-depth comparison showed a total of 3932 DEGs between wild type and *swap1-1* under dark conditions, while under red light 5137 genes were found to be differentially expressed between wild type and *swap1-1* samples (Fig. S9A, Dataset S1, III and V). The heatmap of the top 1000 genes show the gene expression patterns among these samples (Fig. S9B). GO-terms analyses of these comparisons showed significant enrichment of many GO-terms including alternative mRNA splicing, spliceosome mediated splicing, RNA metabolism, photomorphogenesis, red and far-red signal transductions (Dataset S1, II, IV, VI and VIII). Enrichment of these biologically important GO-terms as well as differential expression of several genes involved in light signaling unequivocally highlights the importance of SWAP1 in red light signaling.

### SWAP1 regulates pre-mRNA splicing in Arabidopsis

To identify the extent by which SWAP1 modulates pre-mRNA AS in Arabidopsis, indepth analyses of RNA-seq data were performed using ASpli (Version 2.4.0) (30). To determine the overall differential alternative splicing (DAS) in the mutant, the percentage of inclusion of introns (PIR; Percent Intron Retention) and exons (PSI; Percent Spliced-In) was calculated in four systematic combinations: wild-type_Dark (Col-0_D) vs wild-type_Red light (Col-0_R), *swap1-1 _Dark* (*swap1* _D) vs *swap1-1* _Red light (*swap1* _R), Col-0_D vs *swap1 _D* and Col-0_R vs *swap1*_R. To capture the DAS events with high confidence, only those events which pass through the stringency test of FDR<0.05, fold change≥1.5, and Delta PSI or Delta PIR of ≥ 0.2 were counted. Col-0_D vs *swap1*_D analyses identified a total of 7771 DAS events corresponding to 3957 gene loci, while Col-0_R vs *swap1*_R comparison yielded a total of 12489 DAS events corresponding to 6123 gene loci (Fig. 3A and B; Dataset S2, III and V). While all the subcategories of AS events were found in *swap1-1* samples, the most predominant was the IR event, under both dark and red light-treated conditions (Fig. 3B; Dataset S2, III and V). Interestingly, analysis of overlapping DAS events between dark and red light-treated samples identified a total of 3860 events, corresponding to 50% of dark and 31% of red light events (Fig. 3A), implying a clear shift in SWAP1 targets upon red light illumination.

**Figure 3:**
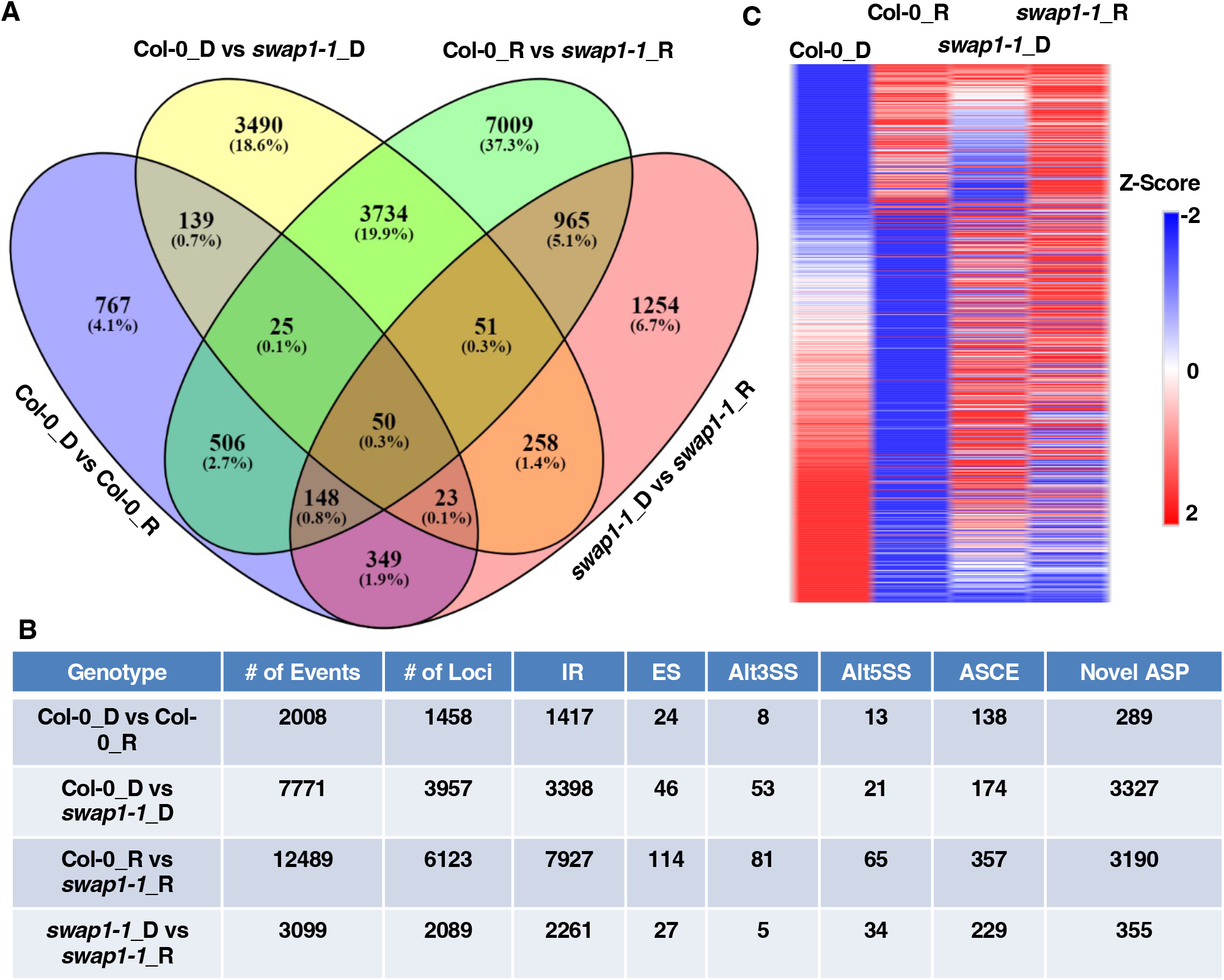
SWAP1 modulates pre-mRNA splicing in Arabidopsis. (A) Venn diagram indicating both number and overlapping differential alternative splicing (DAS) events in wild type and *swap1-1* mutant seedlings under D (Dark) and R (Red light) treated conditions. (B) Table showing the total number of DAS events and the corresponding gene loci numbers detected in analysis. The total number of DAS events is further subdivided into different categories depending on the type of event. (C) Heatmap of top 1000 DAS events plotted based on their Z-scores. Z-scores are calculated based on the basis of their corresponding individual PSI/PIR values under dark and red light treated conditions of wild type and *swap1-1* mutant samples.

Analysis of the scatterplots depicting the splicing efficiency changes among samples indicated that a large number of splicing events show varying degrees of alterations in *swap1-1* compared to wild-type (Fig. S10). Likewise, a heatmap comparison of the overall DAS events also revealed that the splicing patterns of majority of the events in the *swap1-1* mutant are largely opposite to that of wild-type, both under dark and red light-treated conditions (Fig. 3C). The gene list has photoreceptor genes including *phyE* and *PHOT1* (*Phototropin 1*), as well as a number of genes that function downstream of these photoreceptors including *PIFs* (*Phytochrome Interacting Factors*), *COP1* (*Constitutive Photomorphogenic1*), *B-Box* (*BBXs*), and *PAPP2C* (*Phytochrome Associated Protein Phosphatase type 2C*). Furthermore, GO-term analysis showed that a large number of GO-terms are enriched under both dark and light conditions (Dataset S2, IV and VI). GO-terms such as photomorphogenesis, red/far-red light signaling, regulation of flowering, alternative mRNA splicing, and RNA processing were found to be significantly enriched in the dark or red light-treated samples. We also analyzed the number of genes that are affected only at the pre-mRNA splicing or gene expression or both by plotting the Venn diagram of DEGs and DAS. This analysis revealed that a large number of genes were found to be unique to either pre-mRNA splicing or DEGs and only a small number of genes are regulated at both the pre-mRNA splicing and gene expression level (Fig. S11).

To independently verify the RNA-seq data, we randomly selected 5 DAS events for qRT-PCR confirmation. For qRT-PCR, seedlings were grown, treated and RNA was isolated under the same conditions as RNA-seq samples from three independent biological repeats. As shown in Fig. S12, data generated from RT-qPCR is largely similar to that of RNA-seq analysis, implying that the bulk RNA-seq data could independently be reproducible and verifiable.

To investigate the functional relevance of SWAP1 in red light-modulated pre-mRNA splicing, we first determined the DAS events in wild-type (Col-0_D vs Col-0_R) and then compared them against the list of DAS events in *swap1-1* (*swap1* _D vs *swap1*_R). Col-0_D vs Col-0_R analyses identified a total of 2008 DAS events corresponding to 1458 gene loci, while in *swap1-1* mutant a total of 3099 splicing events corresponding to the 2089 gene loci were found to be altered in response to red light (Fig. 3B, Dataset S2, I, VII). Identification of overlapping DAS events interestingly revealed that only 28% and 18% of the overall wild-type and *swap1-1* AS events overlapped, respectively (Fig. 3A), meaning a very large percentage (~82%) of DAS events observed in *swap1-1* are unique. The gene list includes several genes involved in photomorphogenesis and hypocotyl length elongation *(HY2, PIL6,* and *PAPP2*), flowering time regulation (*FCA, VIP4, VEL2*, and *AGLs*) as well as several splicing factor genes including *RS40, PRP31, SR31*, and *SCL30A*. Moreover, a GO-term analysis of unique DAS of *swap1-1* identified several GO-terms related to photomorphogenesis, red/far-red light signaling, pre-mRNA splicing and flowering (Dataset S2, II and VIII). These data clearly indicate that the SWAP1 plays a major role at the molecular level to optimize the global AS of a large number of direct and indirect target genes in response to red light stimulation.

In our previous studies, we have shown that the *ELF3* pre-mRNA is one of the direct targets of both SFPS and RRC1, as both proteins were found to be associated with ELF3 pre-mRNA *in vivo* and regulate its AS pattern (28, 29). In our RNAseq analysis, we identified that *ELF3* is also one of the targets of SWAP1-modulated AS (Fig. S12). We, therefore, presumed that like SFPS and RRC1, SWAP1 might also associate with *ELF3* pre-mRNA. This was tested by performing an RNA-Immunoprecipitation (RIP) followed by a RT-qPCR assay, by employing growth conditions, light treatments, and methods as described previously (28, 29). It was observed that SWAP1 is indeed associated with various regions of *ELF3* pre-mRNA (Fig. S13). This clearly implies that the *ELF3* is one of the direct targets of SWAP1.

### Pre-mRNA splicing of a subset of genes is coordinated by SWAP1, SFPS and RRC1

To investigate the correlation of splicing efficiency changes of light-regulated DAS events among the three splicing mutants, we determined the Delta_PSI/PIR of every individual light-regulated DAS event in wild-type, *sfps-2, rrc1-3*, and *swap1-1*. As shown in the heatmap (Fig. 4A), the splicing efficiency of a majority of light-regulated DAS events was largely similar among *sfps-2, rrc1-3* and *swap1-1* but opposite to that of wild-type, under both dark and red light-treated conditions. A similar pattern of changes in splicing efficiency was also observed in the scatter plots illustrating the wild-type versus individual mutant (WT vs *sfps-2* or *rrc1-3* or *swap1-1*; Fig. S14) or one mutant against another mutant comparisons (*sfps-2* vs *rrc1-3, sfps-2* vs *swap1-1* and *rrc1-3* vs *swap1-1;* Fig. S15)

**Figure 4:**
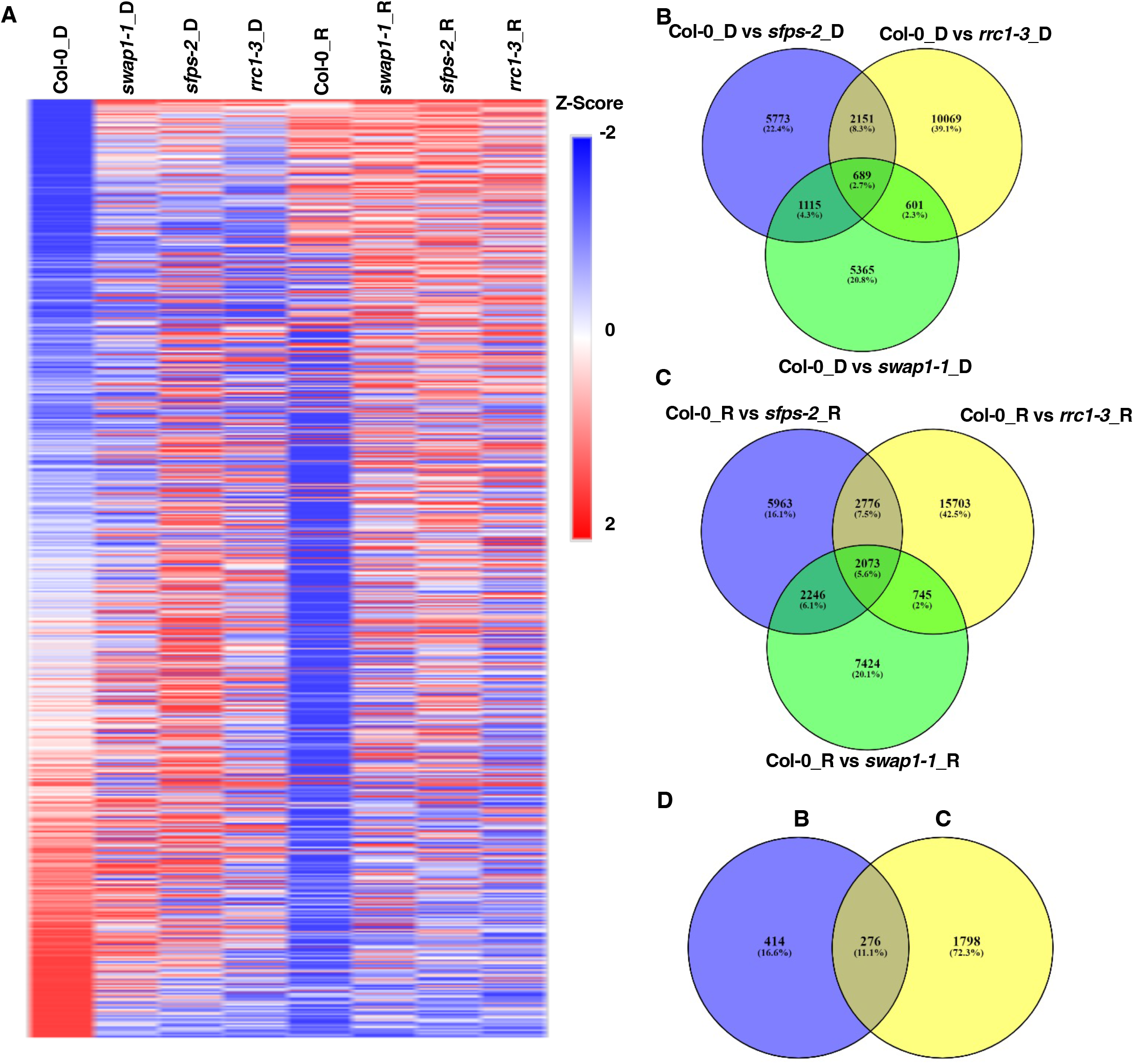
SWAP1, SFPS and RRC1 coordinately regulate pre-mRNA splicing of a subset of genes. (A) Heatmap of top 1000 DAS events plotted based on their Z-scores calculated on the basis of their corresponding individual PSI/PIR values under dark and red light-treated conditions of wild type and different mutant samples. (B) Venn diagram indicating both number and overlapping DAS events in wild type and different mutant seedlings under dark-treated conditions. (C) Venn diagram indicating both number and overlapping DAS events in wild type and different mutant seedlings under red light-treated conditions. (D) Venn diagram showing comprehensive overlapping DAS events observed between (B-dark) and (C-red light) overlapping DAS events.

We have shown above that SFPS, RRC1 and SWAP1 interact with each other forming a ternary complex (Fig. 2C). To test whether these three splicing factors coordinately regulate a subset of common DAS events, a Venn diagram was plotted among the DAS events regulated by each of the splicing factors under dark and red light conditions. Under dark-treated condition, a total of only 689 DAS events were found to be coregulated by all three proteins (Fig. 4A), which accounts for 5.5%, 5.1%, and 8.9% of the total DAS events modulated by SFPS, RRC1 and SWAP1, respectively. Likewise, under red light condition, a total of 2073 DAS events were found to be coregulated by three proteins (Fig. 4B). These accounted for 15.9%, 9.7% and 16.6% of all DAS found to be regulated by SFPS, RRC1 And SWAP1, respectively. When coregulated DAS events from dark and red light-treated samples were overlapped, only 276 DAS events were found to be common between the dark and red light conditions (Fig. 4C). This implies that a very large number of dark and red light-coregulated DAS events are unique to one of the two treatment conditions.

### SWAP1 interacts with red/far-red light photoreceptor phyB

As SWAP1 physically interacts with phyB-interacting SFPS and RRC1, we examined whether SWAP1 also interacts with phyB. Yeast two-hybrid analysis of full-length SWAP1 and phyB showed that these two proteins interact in yeast (Fig. S16). In addition, *in vitro* pull-down assay using bacterially expressed GST-SWAP1 as a bait and full-length phyB-GFP expressed in yeast cells as a prey, showed that GST-SWAP1 interacts preferentially with the phyB-Pfr form (Fig. 5A). GST-PIF1 was used as a control which also interacted strongly with the Pfr form of phyB. To verify the interaction between SWAP1 and phyB, *in vivo* co-immunoprecipitation (Co-IP) assay was performed using *SWAP1_pro_:SWAP1-GFP/swap1-1* transgenic seedlings. Immunoprecipitation of SWAP1-GFP followed by probing for native phyB indicated that SWAP1 does interact with the Pfr form of phyB under *in vivo* condition, albeit in a light-dependent manner (Fig. 5B). As SWAP1 and phyB physically interact with each other and form discrete nuclear speckles and photobodies, respectively, we examined whether the nuclear bodies of these two proteins colocalize. When 5-day-old (Four-days in the dark and one-day in white light) SWAP1-mCherry/phyB-GFP double transgenic seedlings were imaged using confocal fluorescent microscopy, almost all the nuclear speckles of SWAP1-mCherry colocalized with phyB-GFP photobodies (Fig. 5C). Taken together these results clearly show that SWAP1 and phyB colocalize within the nucleus and also physically interact under *in vivo* conditions.

**Figure 5:**
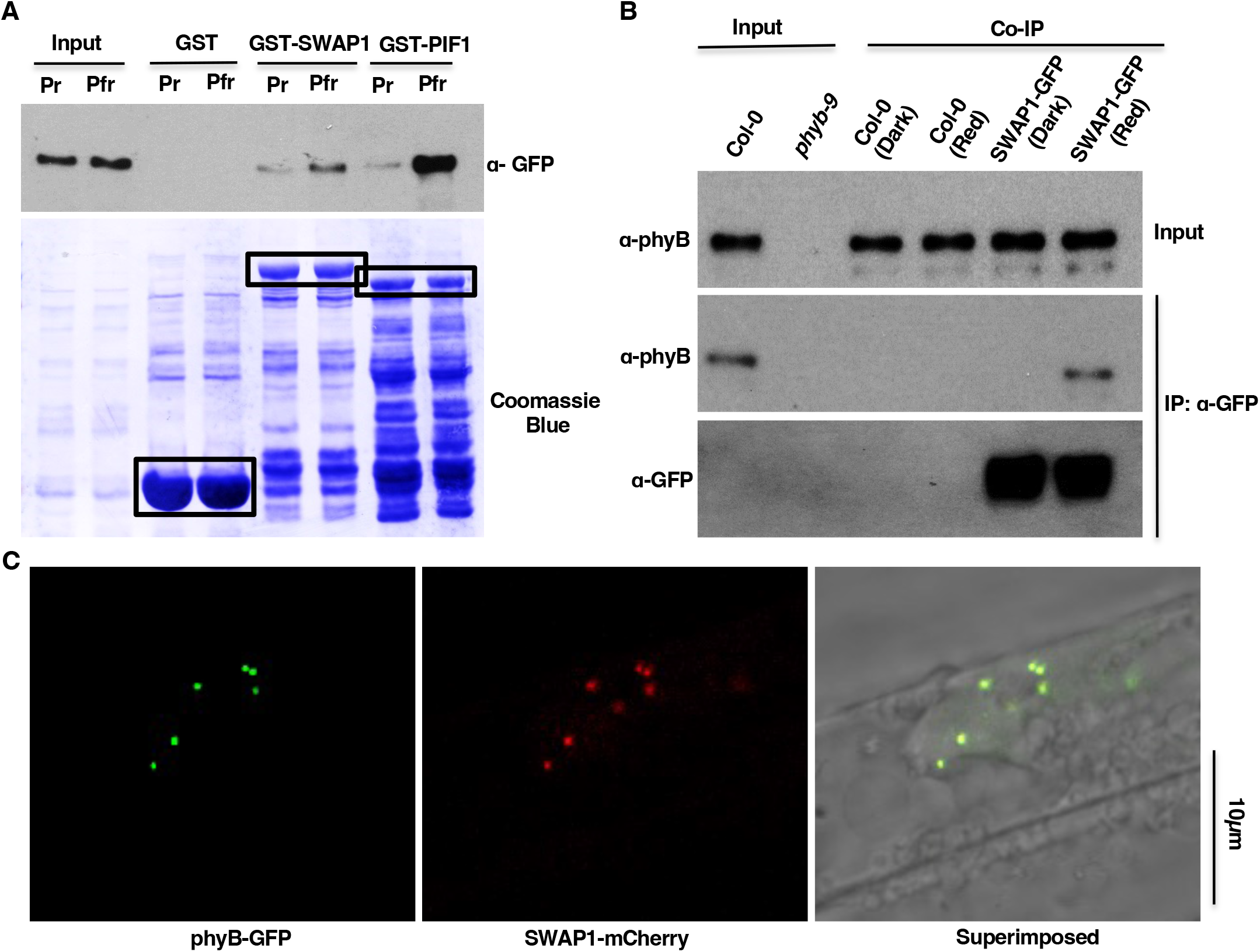
SWAP1 interacts with phyB. (A) Bacterially expressed GST-SWAP1 interacts with phyB-GFP expressed in yeast. Bacterially expressed and glutathione beads bound GST, GST-SWAP1 and GST-PIF1 were mixed with crude extracts of yeast cell-expressed phyB-GFP. One set of tubes was illuminated with red light (Pfr) and another set was exposed to far-red light (Pr) for 10 minutes. Tubes were then incubated for 2hrs at 4°C. After the incubation, beads were pelleted and washed multiple times and proteins were separated on 8% SDS-PAGE. The presence or absence of specific proteins was detected following western blotting with appropriate antibodies. (B) SWAP1 interacts with phyB under *in vivo* conditions. Four-day-old seedlings were either kept in the dark or exposed to the constant red light for 6 hrs (7μmolm^-2^S^-1^) and total protein was extracted in native extraction buffer. 1μg of appropriate antibody bound to Dynabeads (protein A) was added to the protein mixture and incubated for 3hrs at 4°C. After the incubation, beads were pelleted and washed multiple times and proteins were separated on 8% SDS-PAGE. The presence or absence of specific proteins were detected following western blotting with appropriate antibodies. (C)Nuclear speckles of SWAP1-mCherry co-localize with phyB-GFP photobodies. Four-day old dark-grown seedlings of 35Spro:SWAP1-mCherry/35Spro:phyB-GFP double transgenics were exposed to 24hrs of white light and then primary root cells were imaged using confocal microscopy.

## DISCUSSION

Being sessile, plants are routinely exposed to an ever-changing environment. In response to such changes in their surrounding environment, plants deploy diverse strategies, including changes in the growth and developmental pattern(s) to better suit the surrounding conditions. Growth and developmental plasticity in plants (and in other eukaryotes) is often modulated by reprogramming of gene expression globally, which includes pre-mRNA AS and translational regulation (4, 9, 31). The role of AS in transcriptome diversity and gene expression in response to an array of both internal and external stimuli is becoming increasingly evident (4, 9, 10, 22). Light is one such environmental stimuli that shapes plants’ responses by modulating the transcriptome diversity through AS (8, 22, 24). While in the past the core emphasis of the light-mediated signaling was at the level of transcriptional regulation of gene expression, several of the recent deep RNA-seq studies have described the significance of light-mediated changes in AS patterns, transcriptome complexity, and their corresponding effect on plant development (4, 9). Previously, our own study has identified the first-ever known phy interacting splicing factor, SFPS, positively regulating the phy signaling pathway by intricately modulating the pre-mRNA AS (28). Subsequently, Co-IP mass spectrometric study of SFPS-GFP identified numerous proteins including bona fide splicing factors such as RRC1 (29) and SWAP1 (present study).

In this study, we present several lines of molecular, biochemical and genetic evidence to demonstrate that SWAP1 is indeed a phyB interacting bona fide splicing factor, controlling red light-mediated pre-mRNA splicing to regulate photomorphogenesis, in collaboration with SFPS and RRC1. Similar to SFPS and RRC1, SWAP1 protein contains an RNA binding SWAP domain and is categorized as a ‘splicing regulatory’ proteins (Fig. S1C). *swap1* mutant displays red light specific hyposensitivity and flowers early, resembling the *sfps* and *rrc1* mutant phenotypes (Fig. 1; Figs. S2-5) (28, 29). Through a series of *in vitro* and *in vivo* protein-protein interaction assays as well as protein colocalization studies we established that SWAP1 not only colocalizes with SFPS and RRC1 but also interacts with them in a light-independent manner, forming a ternary complex (Fig. 2; Figs. S6 and 7). In addition, SWAP1 interaction with SFPS is not dependent on RRC1. Similarly, SWAP1 interaction with RRC1 also does not require SFPS (Fig. 2). Hence, a single, combination of two proteins and a ternary complex might target a distinct set of pre-mRNAs, in addition to a subset of overlapping targets. This could be verified by identifying the possible unique as well as overlapping direct pre-mRNAs targets of these splicing factors.

One of the characteristics of the splicing factors is their association with the components of one of the U-snRNPs, which enable them to be in proximity to the intron-exon junction of the target (8, 22). SWAP1, like SFPS and RRC1, colocalizes with the components that recognize 3’SS and recruit U2-snRNP to the pre-mRNA branchpoint (Fig. S8) (28, 29). Deep RNA-seq analysis revealed that SWAP1 modulates pre-mRNA AS as well as gene expression, both under dark and red light-treated conditions (Fig. 3; Fig. S9). However, when gene loci related to DAS and DGEs were compared, only a small fraction of them was found to be overlapping (Fig. S11). One possible explanation could be that SWAP1 might first regulate the AS of general splicing and transcription factors, which then in turn modulate the gene expression of downstream genes. Therefore, at least some of the physiological alterations observed in the *swap1* could be due to the combined effect of direct and indirect molecular events.

SWAP1 was found to control pre-mRNA AS of genes not only under red light-treated conditions but also in the dark, albeit ~40% fewer genes than under red light (Fig. 3). However, only a small number of genes were found to overlap between dark and red light-treated conditions. Several light-signaling, hypocotyl length and flowering time modulator genes were found to be selectively spliced differently in response to red light treatment in *swap1-1* (Fig. 2; Dataset S2 III and V). These data clearly imply that the SWAP1 largely targets unique gene loci under two different conditions to modulate optimal plant responses to surrounding ambient light signals. Furthermore, GO analysis of differentially spliced genes revealed the enrichment of GO-terms related to splicing, spliceosomes, RNA processing, red and far-red light signaling, photomorphogenesis, circadian rhythm and regulation of flowering (Dataset S2 IV and VI). As SWAP1, SFPS and RRC1 interact and form a ternary complex, we identified a subset of coregulated splicing events to evaluate the possible functional role of this complex. These data also revealed the subsets of splicing events/genes that are either coregulated by binary interaction between the splicing factors and/or coregulated by the ternary complex in the dark and red light conditions (Fig. 4). Therefore, it is evident that all three proteins target unique pre-mRNAs/genes in addition to a small subset of common targets to cooperatively modulate plant growth and development.

Several phytochrome interacting splicing factors/regulators have been identified in both Arabidopsis (SFPS, RRC1, SWAP1, NOT9B and SWELLMAP2) (24, 26, 28, 29) and *Physcomitrella patens* (PphnRNP-H1, PphnRNP-F1) (32, 33). Since SFPS, RRC1 and SWAP1 also colocalize within phyB photobodies (Fig. 5) (28, 29), it is possible that a ternary complex formed by these proteins might interact with phyB *in planta* to modulate photomorphogenesis via pre-mRNA AS and gene regulation (Fig. S17). This is consistent with recent data showing that the blue light photoreceptor cryptochrome 2 (CRY2) interacts and colocalizes with N^6^-methyladenosine RNA methyltransferase enzymes in CRY2 photobodies (34), implying that photbodies might be sites for mRNA metabolism and pre-mRNA splicing (7). In response to red light perception, phys are known to modulate the protein abundance of numerous downstream interactor proteins (4), however, this does not appear to be the case with any of the three splicing factors (Fig. S3F) (28, 29). Therefore, it is possible that the interaction of these splicing factors with phys might result in some biochemical changes within splicing factors, leading to altered target binding capacity and/or activity. Large-scale proteomic studies have identified multiple phosphorylation sites within SFPS and RRC1 (35, 36). Therefore, it is possible that phys might induce phosphorylation and/or other post-translational modification of these splicing factors within the photobodies, leading to altered target identification and/or protein activity. Alternatively, phys might directly bind to these splicing factors and sequester their activities as has been shown for PIFs (37–39). These possibilities need to be tested in the future to uncover molecular and biochemical mechanisms by which phys/splicing factor complexes regulate pre-mRNA splicing and light-regulated developmental processes.

## MATERIALS AND METHODS

### Plant Materials and growth conditions

All seeds used in this study were in Col-0 background. Seeds were first surface sterilized, plated on MS (without sucrose) medium, and stratified at 4°C for 4-days. Plates were exposed to 3hrs of continuous white light and then transferred to the growth chambers with appropriate light/dark conditions. For hypocotyl length measurement at the seedling stage, digital images of 4-days-old seedlings were obtained, and length was measured by using the ImageJ tool.

Adult plants were grown by transplanting ~10-day-old seedlings to the pots containing Promix soil (Premier Tech Horticulture) and transferred to growth chambers with different light regimes at 22°C. To study flowering time, plants were grown under either long-day (16hrs-light/8hrs-dark) or short-day (8hrs-light/16hrs-dark) conditions at 22°C. Post-bolting rosette leaf number as well as number of days to flower were counted when inflorescence reached ~1cm.

### Vector construction and preparation of transgenic plants

To prepare *SWAP1*_pro_*:SWAP1-GFP*, a genomic DNA fragment containing ~1.6kb promoter region and ~2.7kb gene body was amplified using primers listed in Table S1 and directionally cloned into *pENTR/D-TOPO* vector following the standard protocol (Life Technologies). *pENTR-SWAP1_pro_:SWAP1* vector was recombined into *pGWB4* binary vector. To prepare *35Spro:SWAP1-4XFLAG*, full-length SWAP1 open reading frame (ORF) was amplified using primers listed in Table S1 and cloned into NheI/XmaI restriction sites of pEZS-NL. 4XFLAG DNA fragment was inserted in SmaI/XbaI site of pEZS-NL-SWAP1. SWAP1-4XFLAG was amplified using the primers listed in Table S1 and cloned into SacI/BamHI sites of pCHF1 binary vector. To prepare *35Spro:SWAP1-mCherry*, full-length ORF was amplified using primers listed in Table S1 and cloned into BamHI/XbaI restriction site of the binary vector. All the binary vectors were transformed to GV3101 *Agrobacterium* and then into plants using the floral dip method (40).

To prepare *pVP13-SWAP1* bacterial expression construct, the full-length SWAP1 ORF without the stop codon was amplified using primers listed in Table S1 and directionally cloned into *pENTR/D-TOPO* vector following standard protocol. pENTR-SWAP1 vector was recombined into pVP13 destination vector. The recombined vector was transformed to BL21 (DE3) strain of *E. coli*.

To construct pYES2-SWAP1-GFP, the full-length SWAP1 CDS was amplified using primers listed in Table S1 and cloned into pEZS-NL vector. SWAP1-GFP was digested from pEZS-NL and cloned into pYES2 vector (Life Technologies). pYES2-SWAP1-GFP construct was transformed into RKY1293 *Saccharomyces cerevisiae* yeast cells. To prepare pGAD424-SWAP1 and pGBT9-SWAP1 yeast 2-hybrid vectors, full-length SWAP1 ORF was amplified using primers listed in Table S1 and cloned into EcoRI/BamHI restriction site. These constructs along with their combinations were transformed into yeast strain *AH109*.

### Bacterial and yeast protein induction and purification

To induce the expression of proteins in bacterial culture, ~3ml of well-grown culture at 37°C was transferred to a 500mL flask containing 200mL of LB media and appropriate antibiotics. Flasks were transferred to 37°C rotary shaker for bacterial growth. When the culture reached an OD of ~0.5, isopropyl β-D-thiogalactoside (final concentration 0.1mM) was added and incubated in a rotary shaker for 16 hrs at 18°C. After the incubation, bacteria were first pelleted and then resuspended in protein extraction buffer (100mM Tris-Cl, pH 8.0, 1mM EDTA, pH 8.0, 150mM NaCl, 1X protease inhibitor and 1mM PMSF). The bacterial suspension was sonicated, cell debris was pelleted, and the supernatant was used to purify tagged protein according to standard commercial protocols.

To induce the expression of proteins in yeast culture standard commercial protocol (Thermo Fischer Scientific Inc) was followed, albeit with a small modification. Phycocyanobilin (Frontier Scientific; Cat # P14137) was included in the media during the protein induction. Native protein extraction buffer (29) was supplemented with 1X protease inhibitor and 1mM PMSF. Crude protein extract was stored at −80°C until the assay was performed.

### *In Vitro* pull-down and *In Vivo* Co-IP assays

To perform *in vitro* pull-down assays, GST and MBP tagged proteins were expressed in the bacterial system and purified following standard commercial protocols. An equal quantity of GST-tagged protein was added to the amylose resin-bound MBP or MBP-tagged protein. To investigate the *in vitro* interaction between MBP-SWAP1 and phyB-GFP, an equal quantity of crude extract of phyB-GFP was added to the tubes containing amylose resin-bound MBP or MBP-SWAP1. Dark treatment (phyB-Pr) tubes were constantly kept in the dark, while the red light treatment (phyB-Pfr) tubes were irradiated with the red light (7μmol m^-2^ s^-1^). Sample tubes were incubated in a rotary mixer at 4°C for 3 hrs. After the incubation, beads were pelleted and washed at least 5 times. Following the final wash, beads were immersed in 1X SDS loading buffer, boiled for ~5 minutes, and separated on 6-8% SDS gel. GST and GFP tagged proteins were detected using anti-GST (GE Health care Inc; Cat #RPN 1236) and anti-GFP (Abcam inc: Cat #ab290) antibodies, respectively.

To perform the *in vivo* co-IP assays, 4-day-old dark-grown seedlings were either kept in the dark or irradiated with the constant red light (7μmol m^-2^ s^-1^) for 6 hrs. Sample tissues were frozen and ground thoroughly in native extraction buffer containing appropriate quantities of protease and proteasome inhibitors. Approximately ~1μg of appropriate antibody was added to the protein extract and incubated in a rotary mixer at 4°C for 3 hrs to immunoprecipitate one of the two tagged-proteins. Following the final wash, beads were immersed in 1X SDS loading buffer, boiled at 65°C for ~5 minutes, and separated on 6-8% SDS gel. Interaction between bait and the prey protein was detected using appropriate combinations of primary and secondary antibodies.

### Yeast two-hybrid assays

Different combinations of AD and BD vectors were transformed into yeast strains *AH109/Y187* and multiple positive double transformants were selected on yeast drop-out medium (SD-Ade-Leu-Trp/SD-Leu-Trp). To determine whether the two proteins are interacting within the yeast system, liquid β-galactosidase assay was performed following standard commercial protocol (Clonetech Lab; Matchmaker Two-Hybrid System).

### RNA-seq data analyses

Wild-type and *swap1-1* seedlings were grown under complete darkness for 4 days and then one batch of seedlings in triplicate biological repeats was exposed to continuous red light for 3 hrs (R: 7μmol m^-2^ s^-1^), while the other batch was kept under darkness (D). Following treatment, seedlings were frozen immediately and ground to fine powder. Total RNA was isolated using a commercially available RNA isolation kit following manufacturer’s instructions (Sigma inc: Cat #STRN250-1KT). Total RNA with RIN (RNA Integrity Number) > 6.5 and 28S/18S >1.0 was used for directional library preparation followed by sequencing using NovoSeq PE150 (Novogene, Inc). The sequencing yield is ~200 million reads per sample. For each library, > 90% of the reads were mapped to the unique loci of Arabidopsis TAIR10 genome with the STAR pipeline (41). Differential gene expression was analyzed using DEseq2 (42). Genes with FDR values lower than 0.05 and absolute log two-fold change greater than 0.58 (1.5-fold) were considered as differentially expressed.

AS was analyzed through ASpli (Version 2.4.0) as part of the Bioconductor R package (30), which quantifies the pre-mRNA splicing events through calculating PSI and PIR matrix. In the new version of ASpli, the AS events with an absolute FDR < 5% and Delta PSI_PIR > 20% were deemed differentially spliced. To generate heatmaps and scatterplots, the Z-scores of PSI_PIR values and square roots of PSI_PIR values were calculated as described (29). GO categories belonging to biological processes, molecular functions, and cellular components were analyzed using GeneCodis4. GO terms with P value < 0.05 and FDR <0.05 were considered as significantly enriched.

### RNA extraction and RT-qPCR analysis

To perform RT-qPCR analysis, seedlings were frozen immediately following different treatment conditions. Total RNA was isolated using a commercially available RNA isolation kit following manufacturer’s instructions (Sigma Inc: Cat #STRN250-1KT). Contaminating DNA was removed using RNase-free DNase treatment and 1μg of total RNA was used to synthesize cDNA using reverse transcriptase enzyme following manufacturer’s instructions (Invitrogen: Cat # 28025013). Gene-specific primer sequences used in this study are listed in Table S1. Housekeeping gene *PP2A* was used as an internal control throughout the analyses and the relative expression level was calculated using the comparative C_T_ method.

### RNA Immunoprecipitation assays

Ten-day-old *SWAP1_pro_:SWAP1-GFP/swap1-1* transgenic seedlings grown under 12 hrs light/12 hrs dark conditions were harvested at ZT 12 and immediately cross-linked by vacuum infiltration with 0.5% formaldehyde for 10 mins and then quenched by vacuum infiltration with 0.125 M glycine for 5 mins. Samples were washed with large amounts of autoclaved de-ionized water and dried on filter paper. Samples were immediately frozen and then RNA-IP was conducted following the protocol described previously (28, 29). Primers used in this analysis are listed in Table S1.

## DATA AVAILABILITY

RNA sequencing data were deposited into the Gene Expression Omnibus database (accession number GSE-----). Source data are provided with this paper. Arabidopsis mutants and transgenic lines, as well as plasmids and antibodies generated during the current study are available from the corresponding author upon reasonable request.

## ACKNOWLEDGEMENTS

We thank members of the Huq laboratory for critical reading of the manuscript. This work was supported by grants from the National Science Foundation (MCB-2014408) to E.H. and (MCB-2014542) to A.S.N.R. The authors acknowledge the Texas Advanced Computing Center (TACC) at The University of Texas at Austin for providing High Performance Computing, visualization, and database resources that have contributed to the research results reported in this paper.

## Notes

### Competing Interest Statement

The authors have declared no competing interest.

