## Supplemental figures for "SWAP1-SFPS-RRC1 splicing factor complex modulates pre-mRNA splicing to promote photomorphogenesis in Arabidopsis"

### SUPPLEMENTARY FIGURES

**A**

| Accession | Spectrum Number (Control) | Spectrum Number (SFPS) | Unique Peptides | Protein Coverage |
|---|---|---|---|---|
| At1g30480 (SFPS) | 0 | 872 | 25 | 0.3979 |
| At4g31200 (SWAP1) | 0 | 31 | 12 | 0.2446 |

**B**

1 MDRRQHDYAA SSGLPYAQQQ QQQGPNFQQQ QQPQFGFHPQ HPQYPSPMNA
51 SGFIPPHPSM QQFPYQHPMH QQQQPQHLPH PPHPQMFGQQ QPQAFLPHLP
101 PHHLPPPFPG PYDSAPPPPP PADPELQKRI DKLVEYSVKN GPEFEAMMRD
151 RQKDNPDYAF LFGGEGHGYY RYKHFLSMHP PGGPFDPPFP SSSMPMIHHP
201 PNPMMSPSMN NVPGALAVPP IRQPPFPPFH DHHQLQQHLP QPHPFAPHAR
251 PDFDQSTHAF RGLSGPLPAD VAMELNGVLG NLNGTKESIK SAKIWFMQRS
301 PFAPALAEAL RDRVFAMDDS DRQMHIVYLA NDILFDSLQR RTNLHEFDNE
351 ALAFRPILGS MLGRIYHFPQ NKEENQSRLE KILQFWASKE VFDQDTISSL
401 EKEMKSGPPA NTFSHSPIIA AHALQRPGML QQPPNSNVSS TMNLEHLTNP
451 VATQQFIPNV MPPGAFPGSI PLNASVPPPT QPPAGEKPPP YPLFPPGLIP
501 GMVRKMQIGS GVPYSPLSPL DIPTVIPPSD TPQSEVLERV SKFFKEIGEV
551 NPSEGPMGSE SQDDYDNYER DSPQRKGGAC IPPPPNLQVD PETGTYADGS
601 TDKKSGSGRL GLGATADPNE PTQYDDVYTS YRKHRSTNYH TSMSARATTR

**C**

**At4g31200 (SWAP1)**

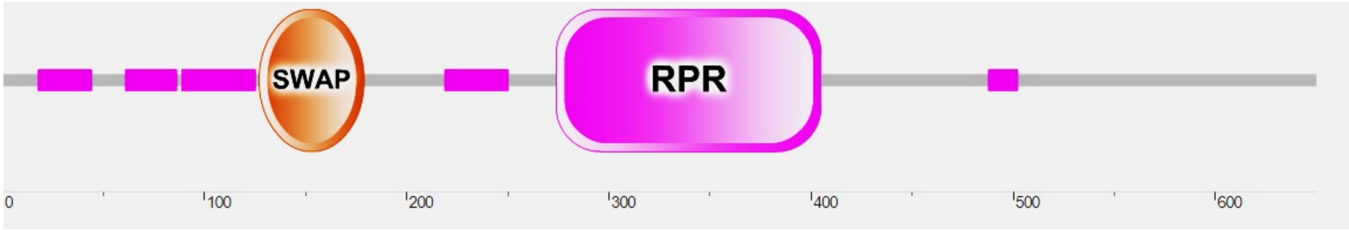


**Fig. S1: Identification of SWAP1 as SFPS interactor in IP-MS (Immunoprecipitation followed by mass spectrometry) assay.**

(A) SWAP1 co-immunoprecipitates with SFPS-GFP. Four-day-old dark-grown $_{SFPSpro}$:*SFPS-GFP/sfps-2* seedlings were either kept in the dark or irradiated with red light (7µmol $m^{-2}$ $s^{-1}$) for 6 hrs. SFPS-GFP was immunoprecipitated using anti-GFP antibodies and interacting proteins were identified through MS-MS. (B) SWAP1 protein sequence showing peptides (colored) detected in MS-MS. (C) Predicted conserved domains of SWAP1 protein.

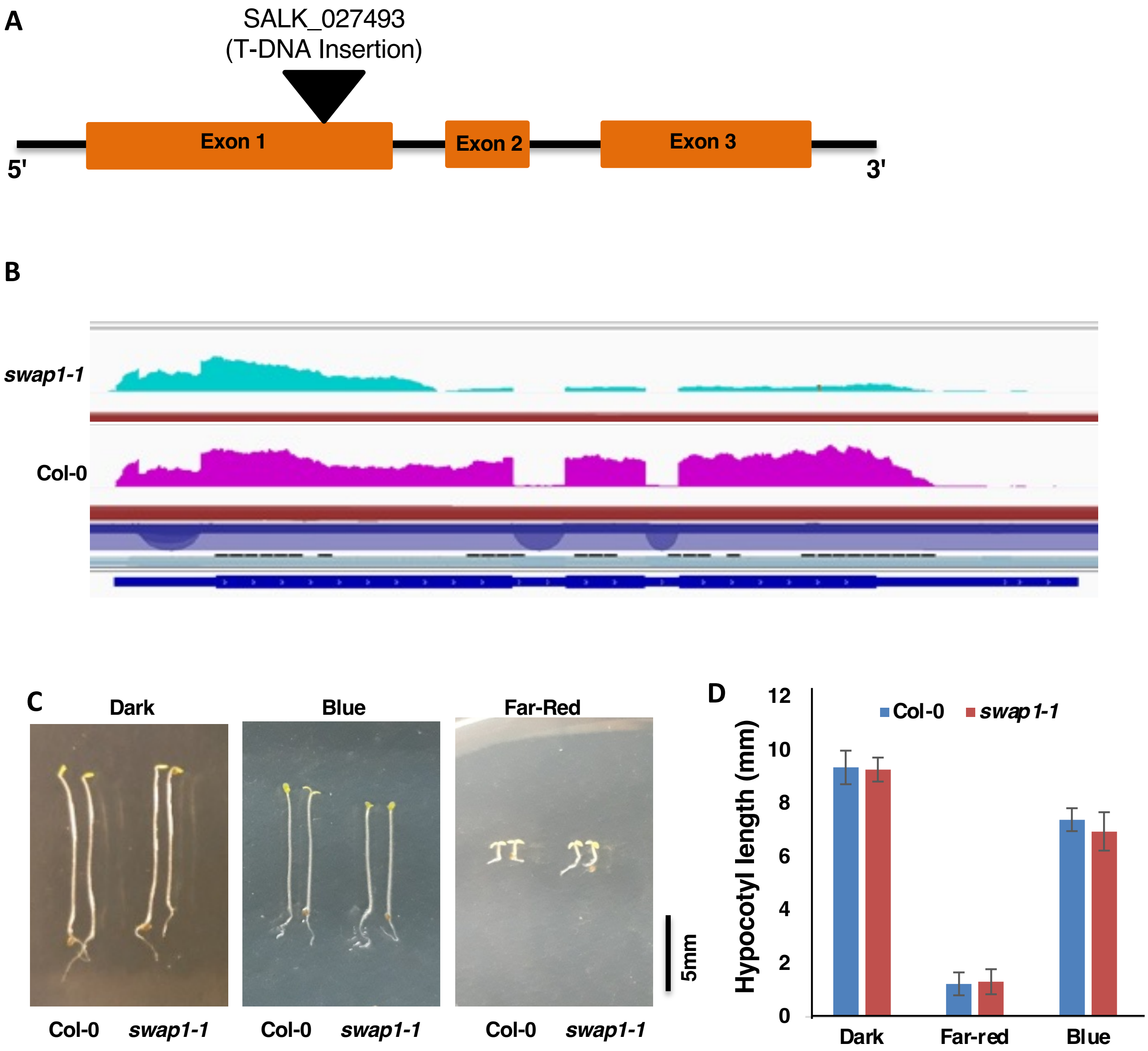


**Fig. S2: *swap1-1* mutant seedlings display wild-type sensitivity on blue and far-red light**
(A). Schematic diagram showing the gene structure of SWAP1 and the T-DNA insertion site of *swap1-1* (SALK_027493). Lines are not to the scale. (B) IGV visualization of *SWAP1* expression in Col-0 and *swap1-1* mutant. (C) Digital images of representative seedlings of different genotypes grown under either continuous dark or continuous far-red-light or blue-light for 4 days. (D) Quantification of the hypocotyl length of 4-day-old seedlings grown under either continuous dark or continuous far-red light or blue light. Bars indicate mean length in mm and error bars indicate standard deviation. Statistical significance between Col-0 and *swap1-1* was determined by student's T-Test.

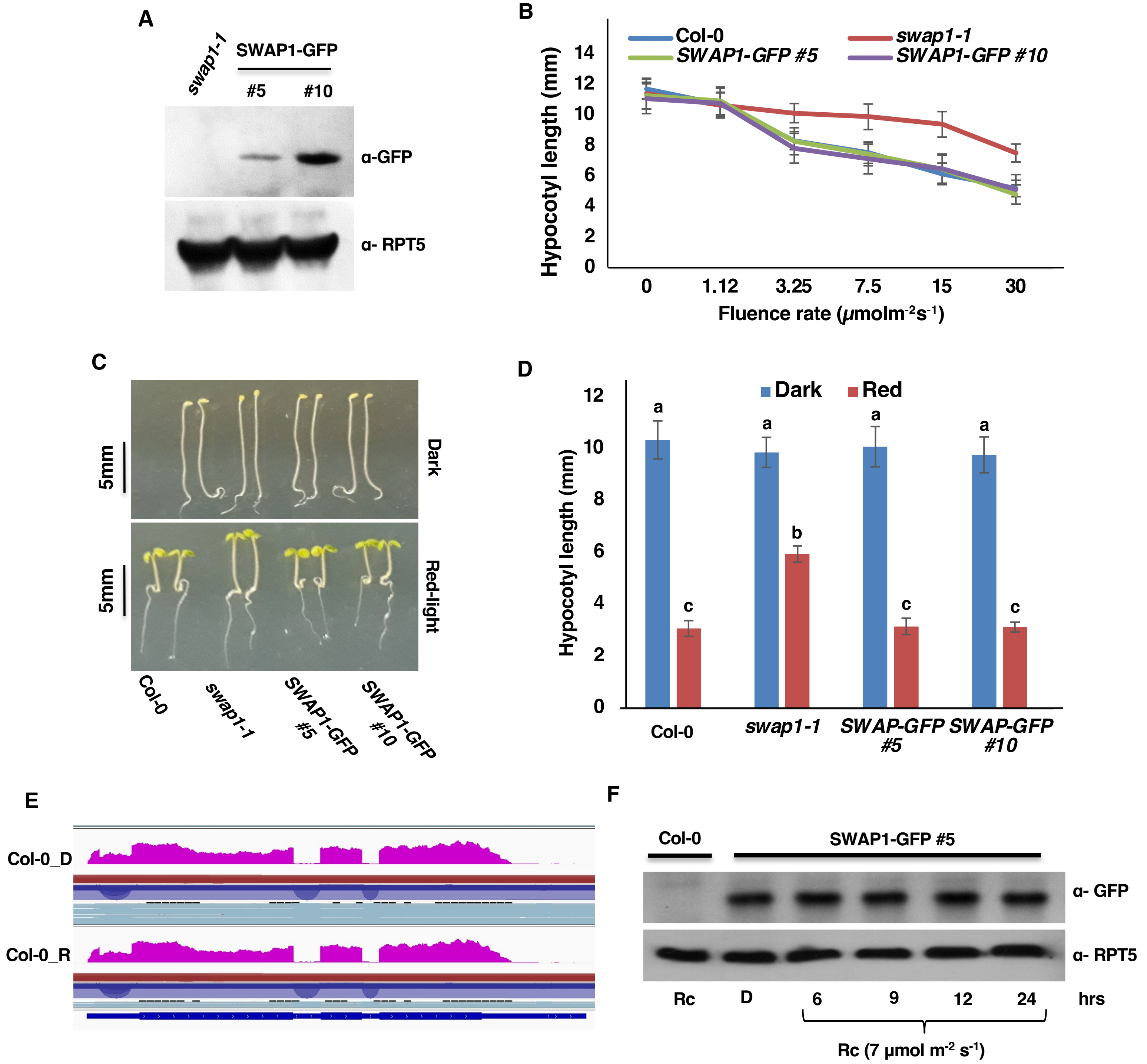


**Fig. S3: *SWAP1$_{pro}$:SWAP1-GFP* complements the *swap1-1* mutant phenotype under red light**
(A) SWAP1-GFP protein quantification in two independent homozygous *SWAP1$_{pro}$:SWAP1-GFP* transgenic seedlings. (B) Quantification of the hypocotyl length of 4-day-old seedlings of two independent transgenic lines expressing SWAP1-GFP under its own promoter along with *swap1-1* grown under either continuous dark or continuous red light of different fluence rate. (C and D) Digital images of representative seedlings (B) and bar graph showing hypocotyl lengths (C) of different genotypes grown under either continuous dark or continuous red light (7µmolm$^{-2}$s$^{-1}$) for 4 days. Bars indicate mean length in mm and error bars indicate standard deviation. Statistical significance among different genotypes was determined using single factor ANOVA followed by Tukey's post hoc analysis and is indicated by different letters. (E) ) IGV visualization of *SWAP1* expression in Col-0 under dark and red-light irradiated conditions. (F) SWAP1-GFP protein abundance in dark and red light treated seedlings. Total protein was extracted in denaturing buffer and protein was separated on 10% SDS-PAGE gel. Protein was transferred to a PVDF membrane and the presence of SWAP1-GFP was detected using α-GFP antibody. RPT5 protein was used as an internal control.

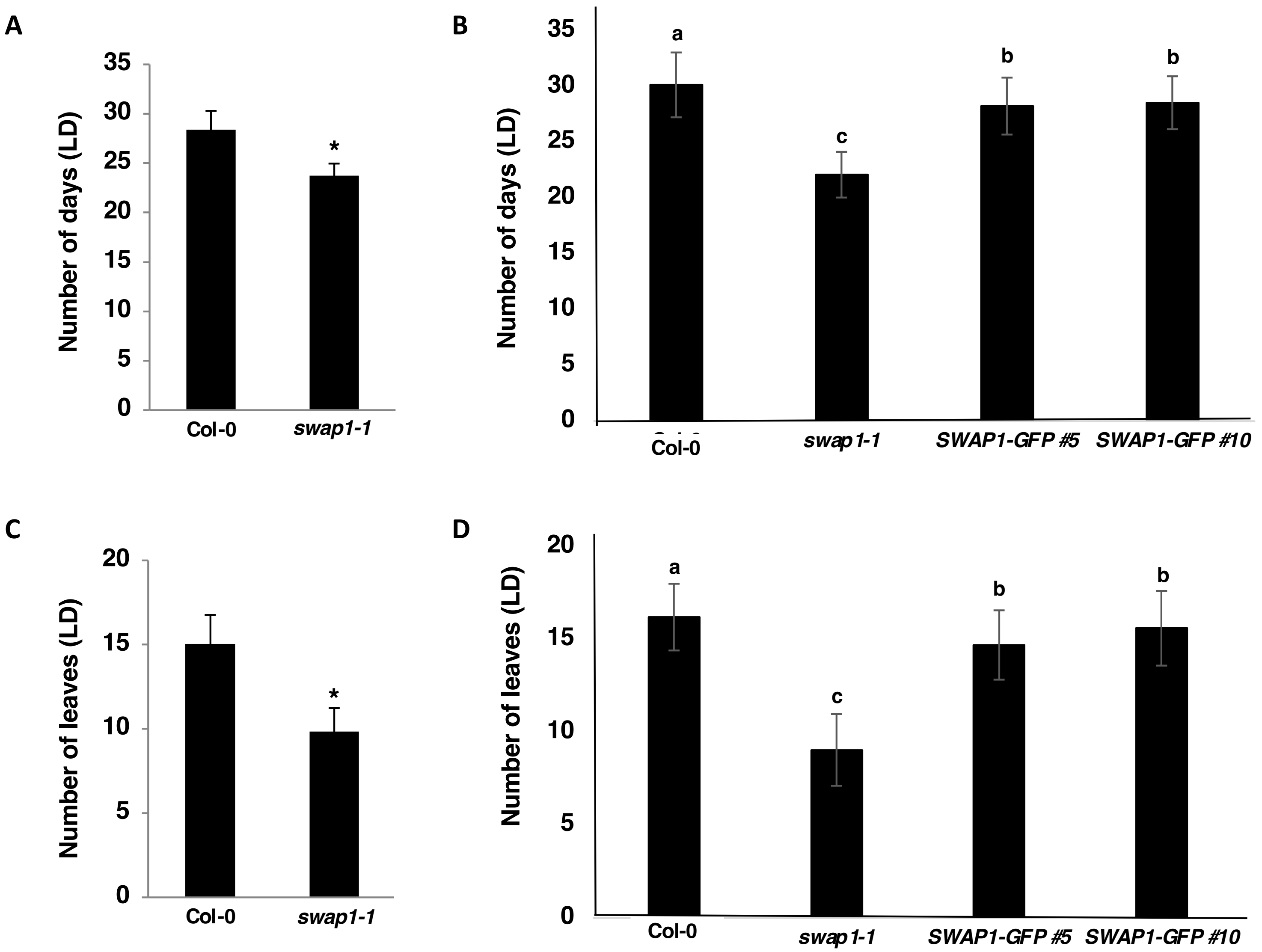


**Fig. S4:** ***swap1-1*** **mutant plants flower early under Long-day conditions.**

(A) And (B) Number of days it takes plants show 1cm bolting under long-days (LD: 16hrs light/8hrs dark) condition. (C) and (D) Number of rosette leaves plants have when plants show 1cm bolting under LD condition. Each data point represents the mean average of at least 20 plants and the error bars show SD. Either Student's T-Test or one-way ANOVA followed by Tukey's post-hoc test was carried out to determine the statistical significance between or among samples, respectively.

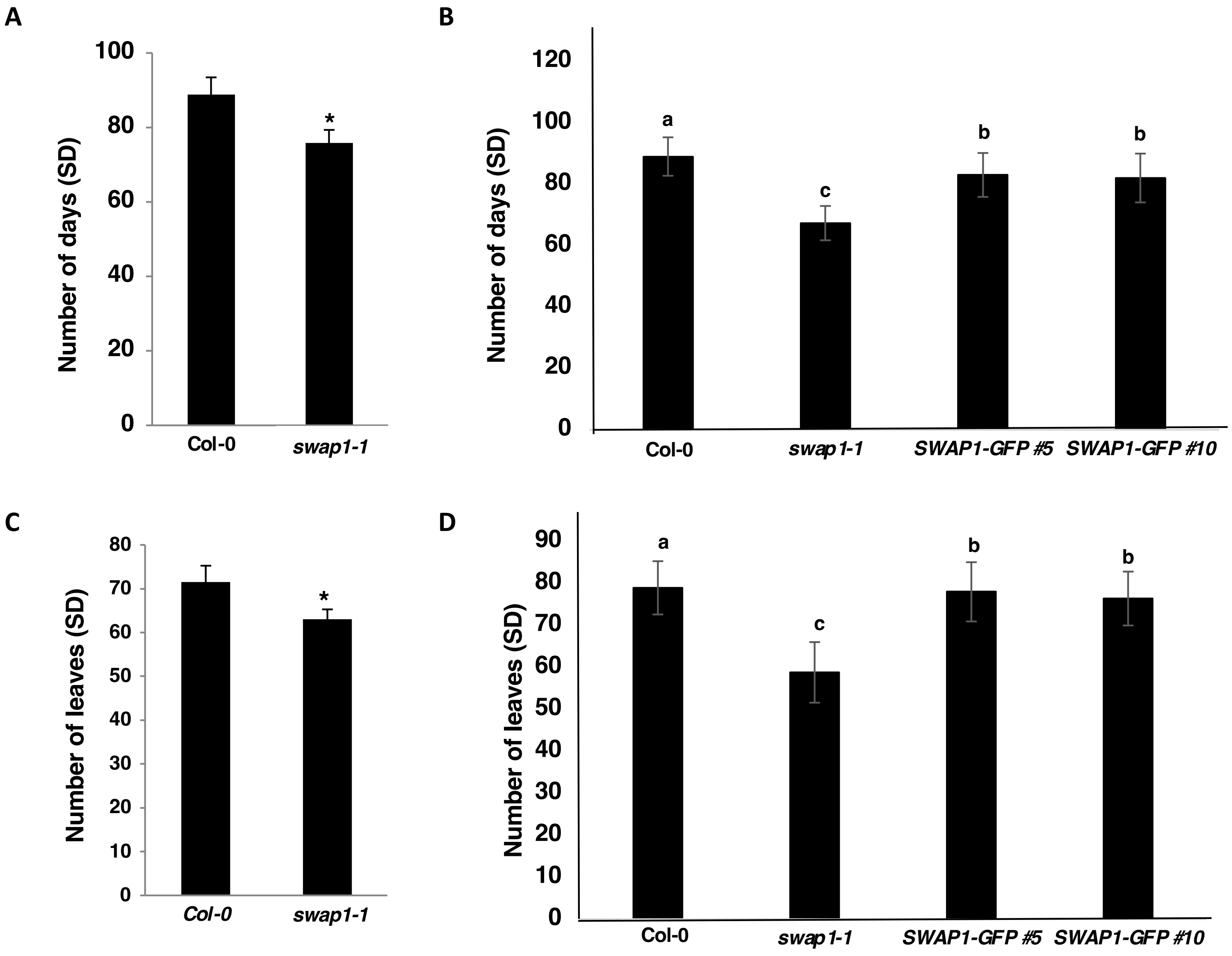


**Fig. S5:** ***swap1-1*** **mutant plants flower early under short-day conditions.**

(A) And (B) Number of days it takes plants show 1cm bolting under long-days (SD: 8hrs light/16hrs dark) condition. (C) and (D) Number of rosette leaves plants have when plants show 1cm bolting under LD condition. Each data point represents the mean average of at least 20 plants and the error bars show SD. Either Student's T-Test or one-way ANOVA followed by Tukey's post-hoc test was carried out to determine the statistical significance between or among samples, respectively.

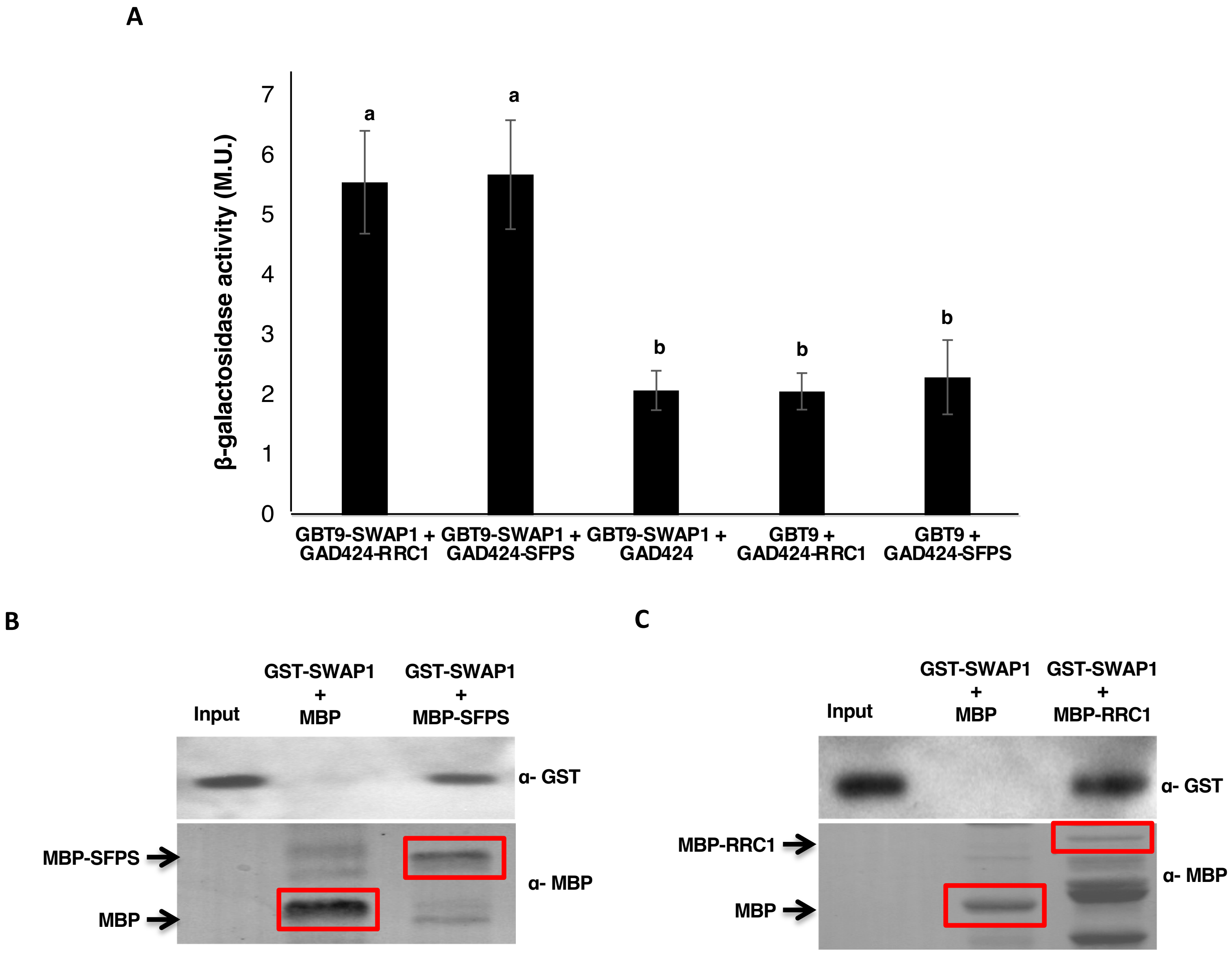


**Fig. S6: SWAP1 interacts with SFPS and RRC1**

(A) SWAP1 interacts with both SFPS and RRC1 in a yeast 2-hybrid system. Full-length SWAP1 CDS was inserted in pGBT9 vector, while full-length CDS of SFPS and RRC1 were inserted in pGAD424 vector. Liquid β- Galactosidase activity assay was carried out to quantify the strength of interaction between SWAP1 and SFPS/RRC1. Each data point reflects the average of three values and the error bars represents the SD. One-way ANOVA with Tukey's post-hoc test ($P<0.01$) was carried out to determine the statistical significance among the samples.

(B) GST-SWAP interacts with MBP-SFPS and MBP-RRC1 (C) *in vitro*. Bacterially expressed and amylose resin-bound MBP, MBP-SFPS and MBP-RRC1 were used as bait proteins to pull-down GST-SWAP1 prey protein. The pull-down reaction was incubated for 60mins at $4^0$C on a rotating shaker. After the incubation, beads were washed at least 5 times. Proteins were separated on 10%SDS-PAGE gel, transferred to PVDF membrane and first immunoblotted with α- GST, followed by α- MBP.

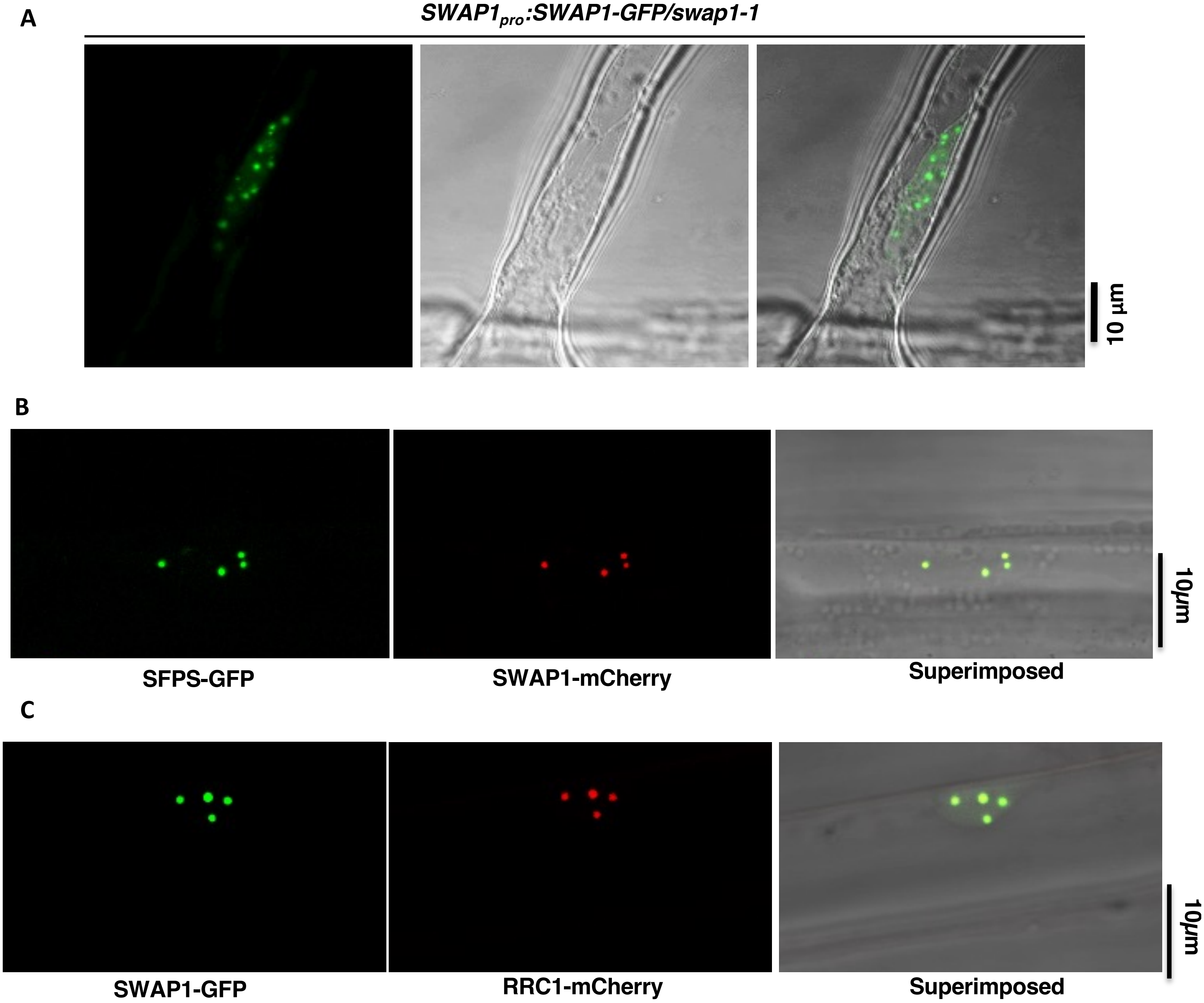


**Fig. S7: SWAP1 nuclear speckles col-localize with SFPS and RRC1**
(A) SWAP1-GFP forms nuclear speckles. (B) SFPS-GFP and SWAP1-mCherry nuclear speckles co-localize with each other. (C) SWAP1-GFP and RRC1-mCherry nuclear speckles co-localize with each other.

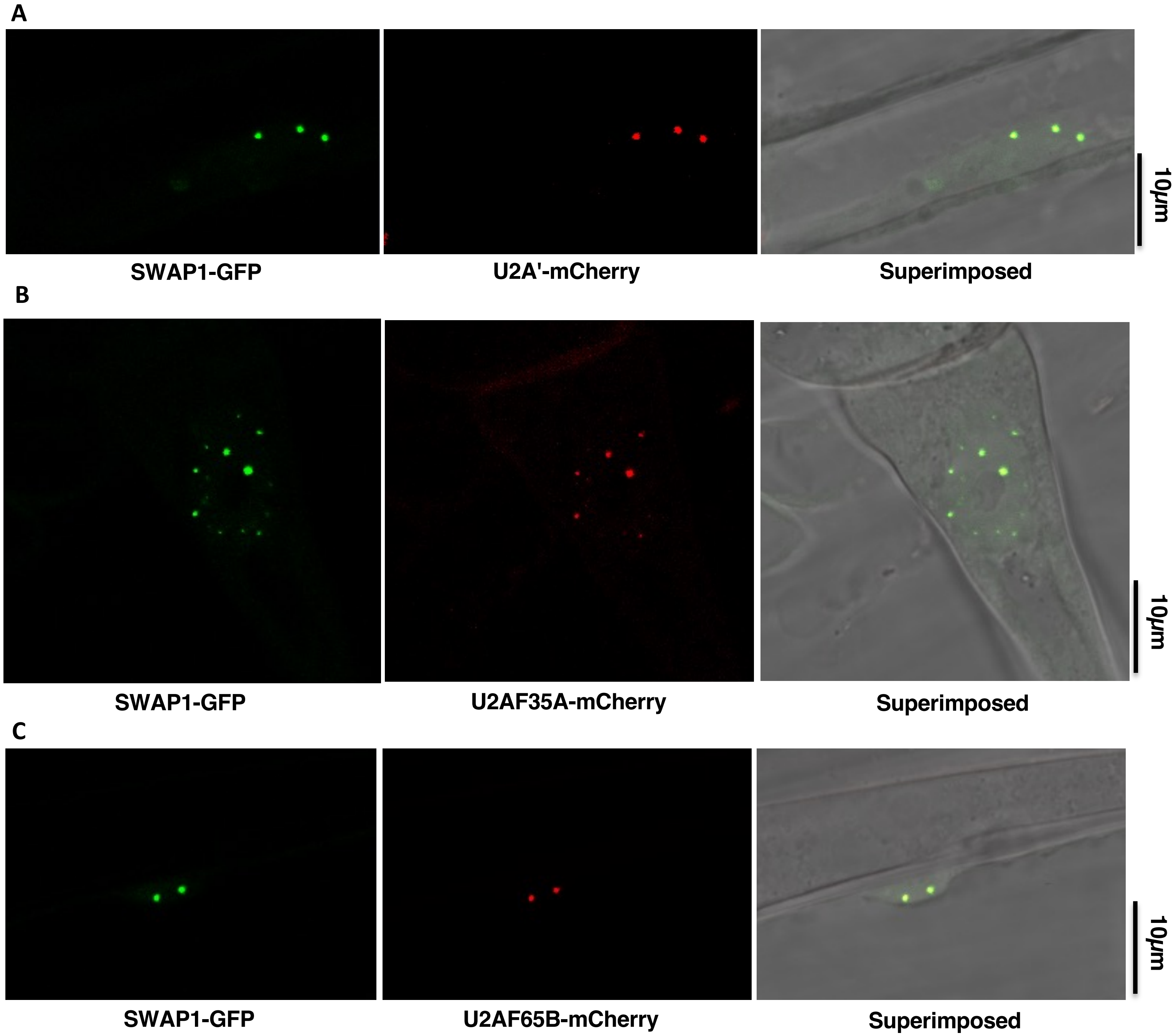


**Fig. S8: SWAP1 nuclear speckles col-localize with U2- snRNPs associated components**
(A) SWAP1-GFP nuclear speckles co-localize with nuclear speckles of U2A'-mCherry. (B) SWAP1-GFP nuclear speckles co-localize with nuclear speckles of U2AF35A-mCherry. (C) SWAP1-GFP nuclear speckles co-localize with nuclear speckles of U2AF65B-mCherry.

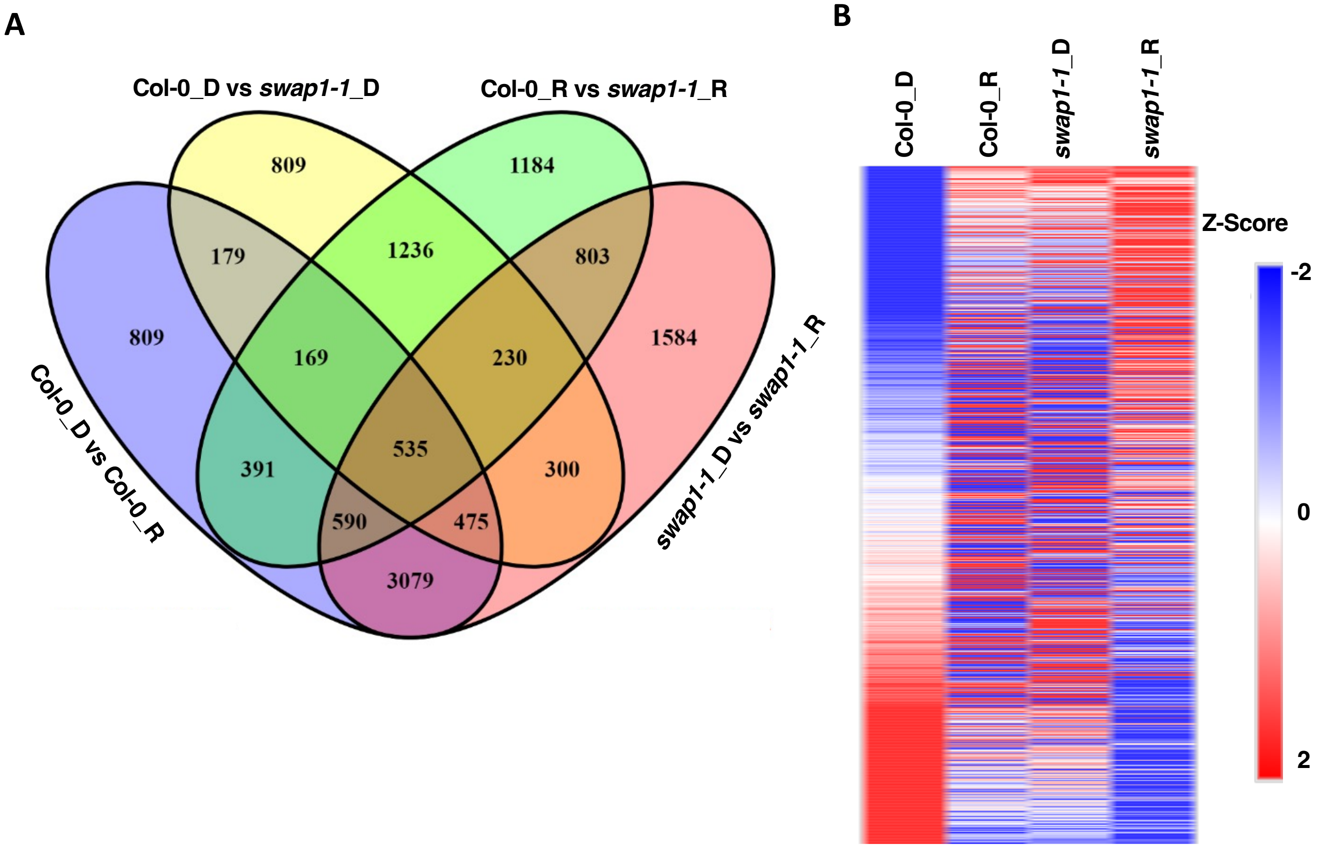


**Fig. S9: SWAP1 regulates expression of a large number of genes both in dark and light conditions**
(A) Venn diagram indicating both number and overlapping differentially expressed genes (DEGs) events in wild type and *swap1-1* mutant seedlings under D (Dark) and R (Red light) treated conditions.
(B) Heatmap of top 1000 DEGs events plotted based on their Z- scores. Z- scores are calculated based on the basis of their corresponding individual expression values under dark and red light treated conditions of wild type and *swap1-1* mutant samples.

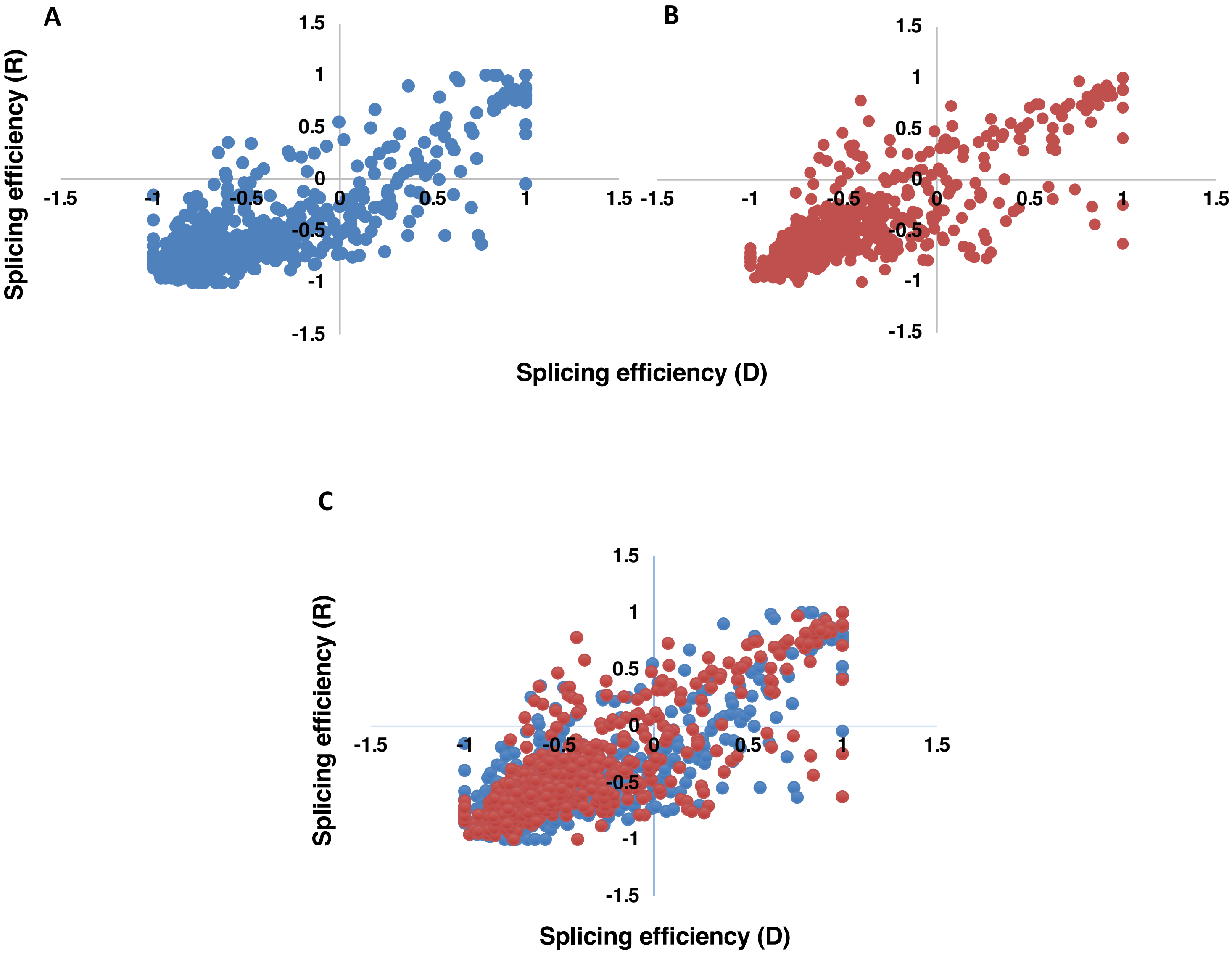


**Fig. S10: *swap1-1* exhibits altered splicing efficiency compared to wild-type**
Scatter plots exhibiting the splicing efficiency changes in wild-type (A) and *swap1-1* (B) mutant samples. (C) Overlapping of splicing efficiency observed in (A) and (B). The X-axis represents the splicing efficiency of samples in the dark, while Y-axis represents the splicing efficiency of samples under red light.

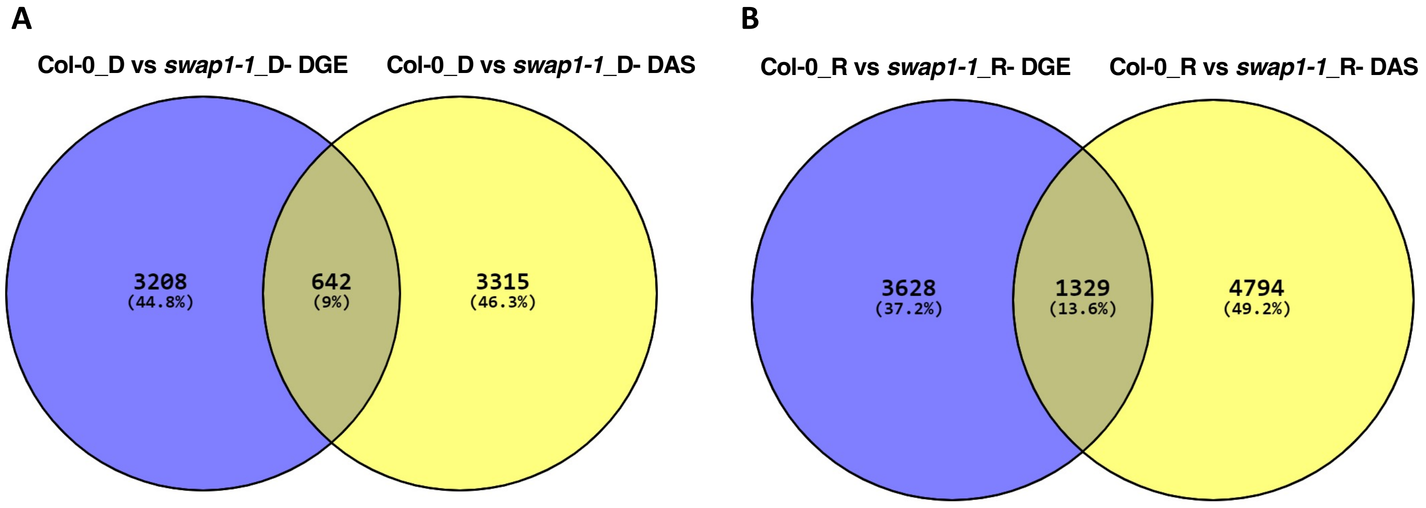


**Fig. S11: Overlap of DEGs and DAS genes modulated by SWAP1**

Venn diagram indicating both number and overlapping of DEGs and DAS genes in comparison of wild type vs *swap1-1* in the dark (A) and red light (B) illuminated conditions.

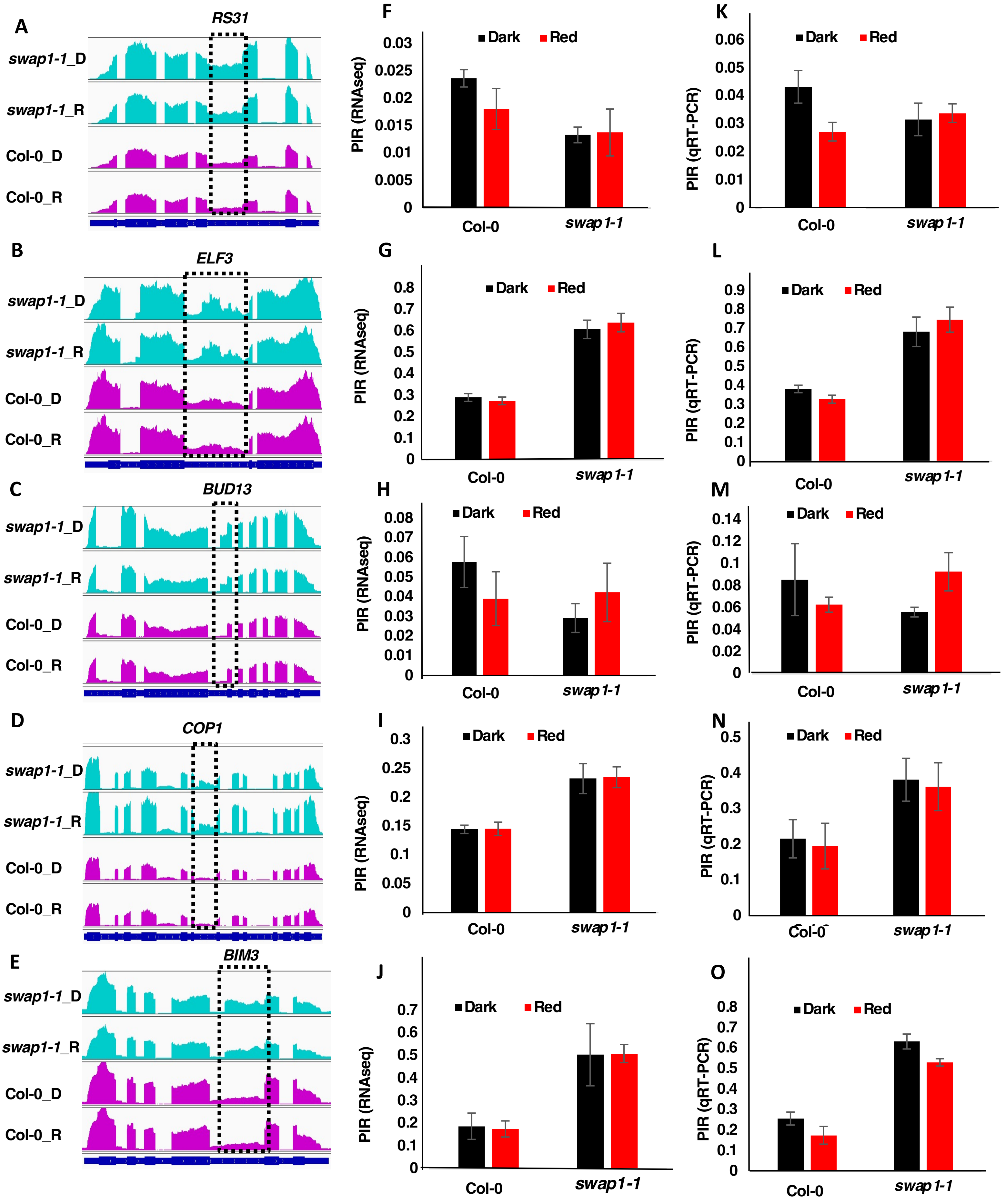


Fig. S12: qRT-PCR confirmation of RNAseq data

**Fig. S12: qRT-PCR confirmation of RNAseq data**

(A-E) IGV visualization of read depth graphs depicting the DAS events in (A) *RS31,* (B) *ELF3,* (C) *BUD13,* (D) *COP1* and (E) *BIM3* as detected from RNA-seq analysis of Col-0 vs *swap1-1* samples. (F-J) Intron retention patterns of corresponding genes as obtained from RNA-seq analysis of Col-0 vs *swap1-1* samples. (H-O) Quantification of retained introns of the corresponding genes as determined independently by qRT-PCR analysis.

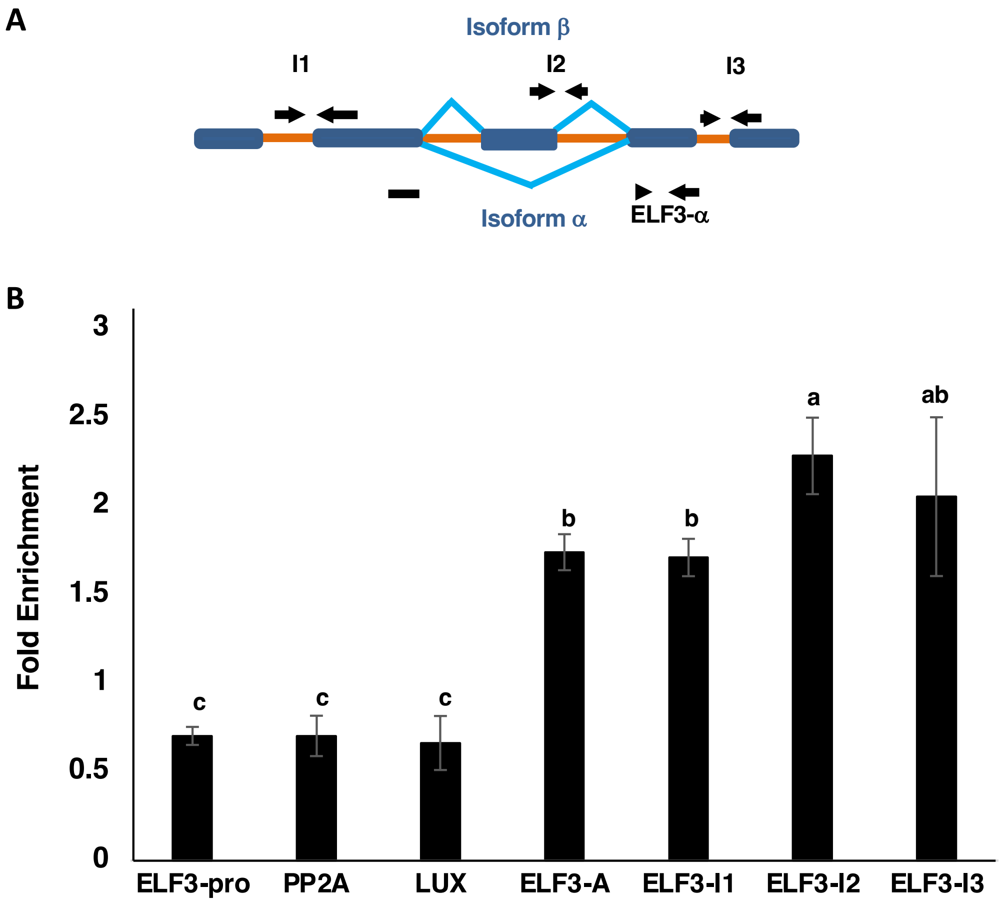


**Fig. S13: SWAP1 associates with *ELF3* pre-mRNA under *in vivo* condition**

(A) Schematic diagram showing gene structure, introns and exons, different isoforms as well as primer positions used in the RNA Immunoprecipitation-qRT-PCR assay. (B) Fold enrichment of different parts of ELF3 pre-mRNA following immunoprecipitation of SWAP1-GFP indicates that SWAP1 associates with ELF3 pre-mRNA under *in vivo* conditions. RNA/SWAP1-GFP complex was immunoprecipitated (IP) and extracted using anti-GFP antibodies. RNA was carefully purified from IP samples and reverse transcribed into cDNA following standard protocol. The abundance of each region in the transcript was quantified by qRT-PCR method. Each bar is the mean ± SEM (n= 3 biological repeats). Statistical difference was calculated based on One-way ANOVA followed by Tukey's post hoc analysis. Different letters indicate statistical significance among the gene parts.

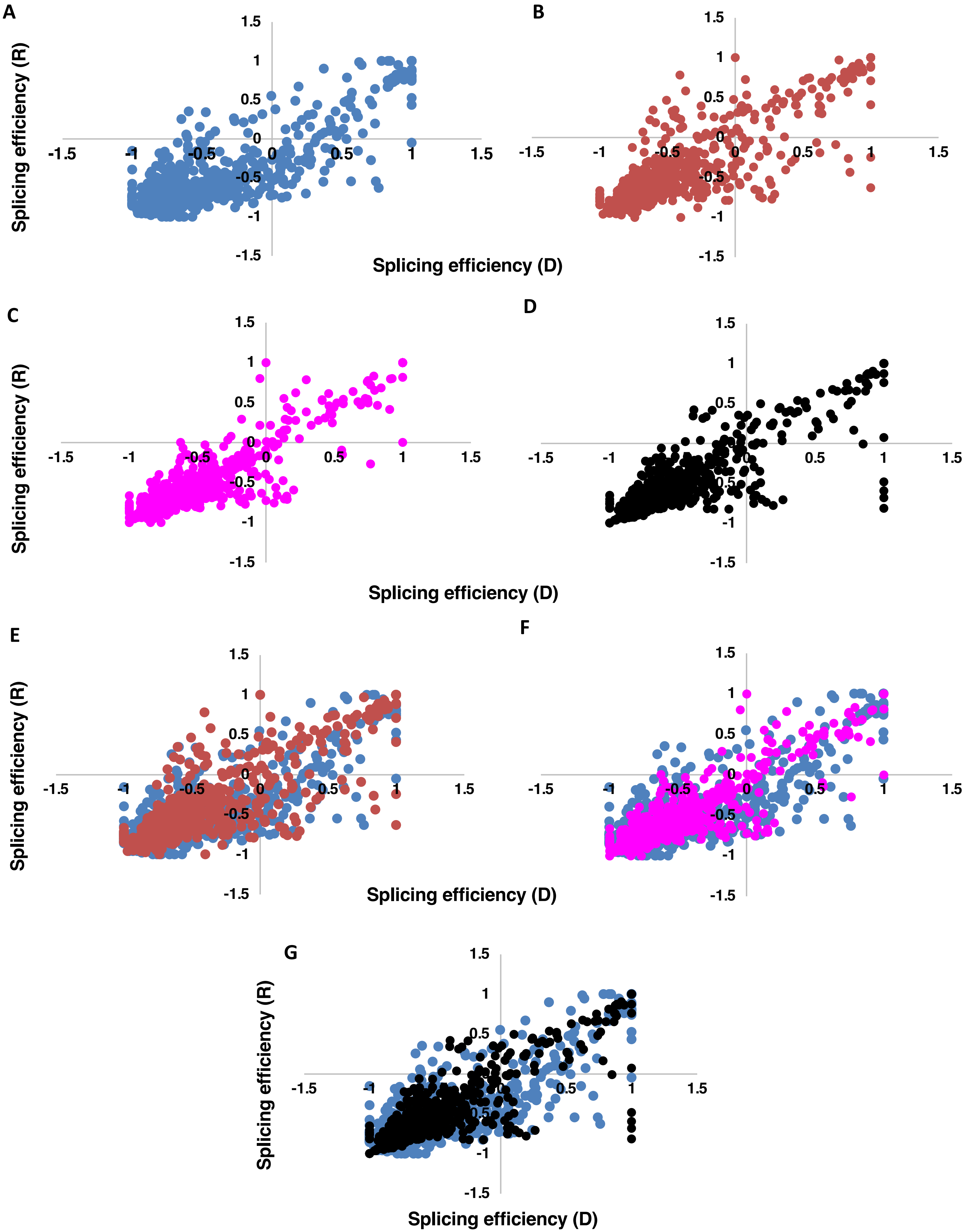


**Fig. S14:** ***sfps-2*, *rrc1-1* and *swap1-1* exhibit altered splicing efficiency compared to wild-type**

**Fig. S14: *sfps-2*, *rrc1-1* and *swap1-1* exhibit altered splicing efficiency compared to wild-type**
Scatter plots exhibit the splicing efficiency changes in the wild-type (A), *swap1-1* (B), *sfps-2* (C) and *rrc1-3* (D) mutant samples. (E) Overlapping of splicing efficiency observed in (A) and (B). (F) Overlapping of splicing efficiency observed in (A) and (C). (G) Overlapping of splicing efficiency observed in (A) and (D).
The X-axis represents the splicing efficiency of samples in the dark, while Y-axis represents the splicing efficiency of samples under red light.

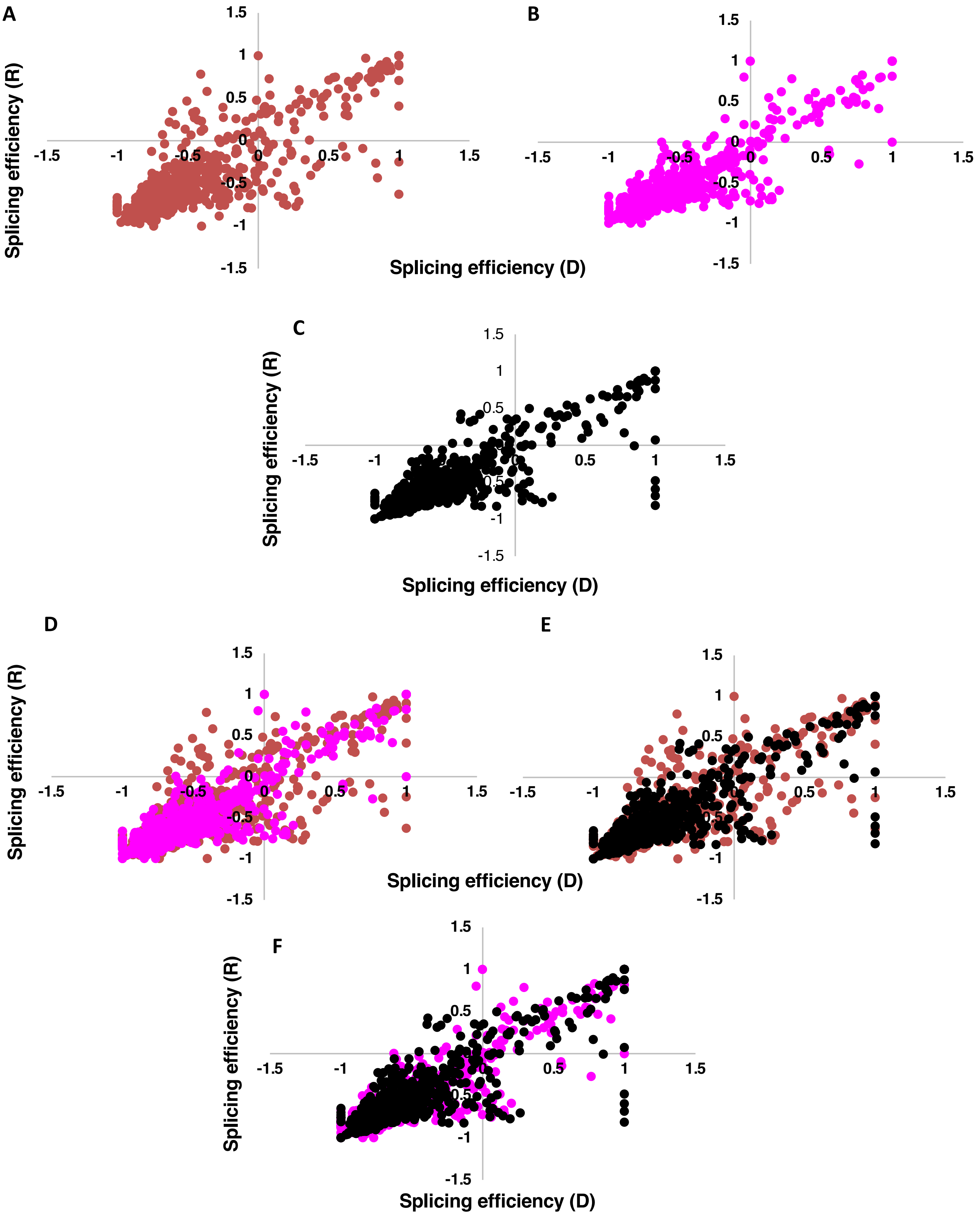


**Fig. S15: The light-regulated splicing efficiency is largely similar in all three mutants**

**Fig. S15: The light-regulated splicing efficiency is largely similar in all three mutants**

Scatter plots exhibit the splicing efficiency changes in *swap1-1* (A), *sfps-2* (B) and *rrc1-3* (C) mutant samples. (D) Overlapping of splicing efficiency observed in (A) and (B). (E) Overlapping of splicing efficiency observed in (A) and (C). (F) Overlapping of splicing efficiency observed in (C) and (D). The X-axis represents the splicing efficiency in the dark samples, while Y-axis represents the splicing efficiency under red light samples.

A

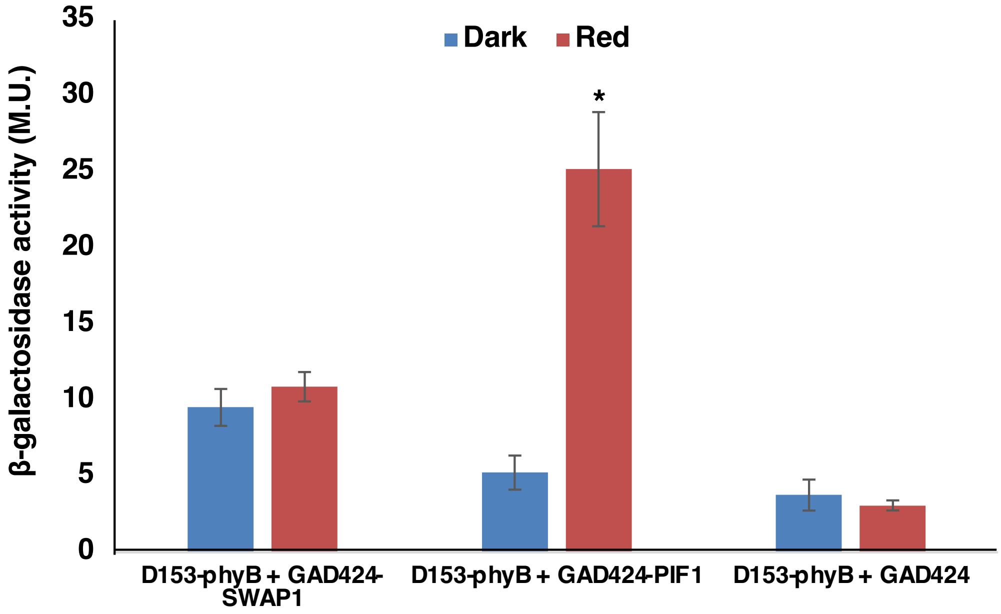


**Fig. S16: SWAP1 interacts with phyB in yeast 2-hybrid system**

Full-length *SWAP1* CDS was inserted in vector containing the GAL4 activation domain (AD), while the full-length *phyB* CDS was fused to the GAL4 DNA binding domain (BD). Liquid β- Galactosidase activity assay was carried out to quantify the strength of interaction between SWAP1 and phyB. Each data point reflects the average of three values and bars represent the SD. Student's T-Test was carried out to determine the statistical significance among the samples.

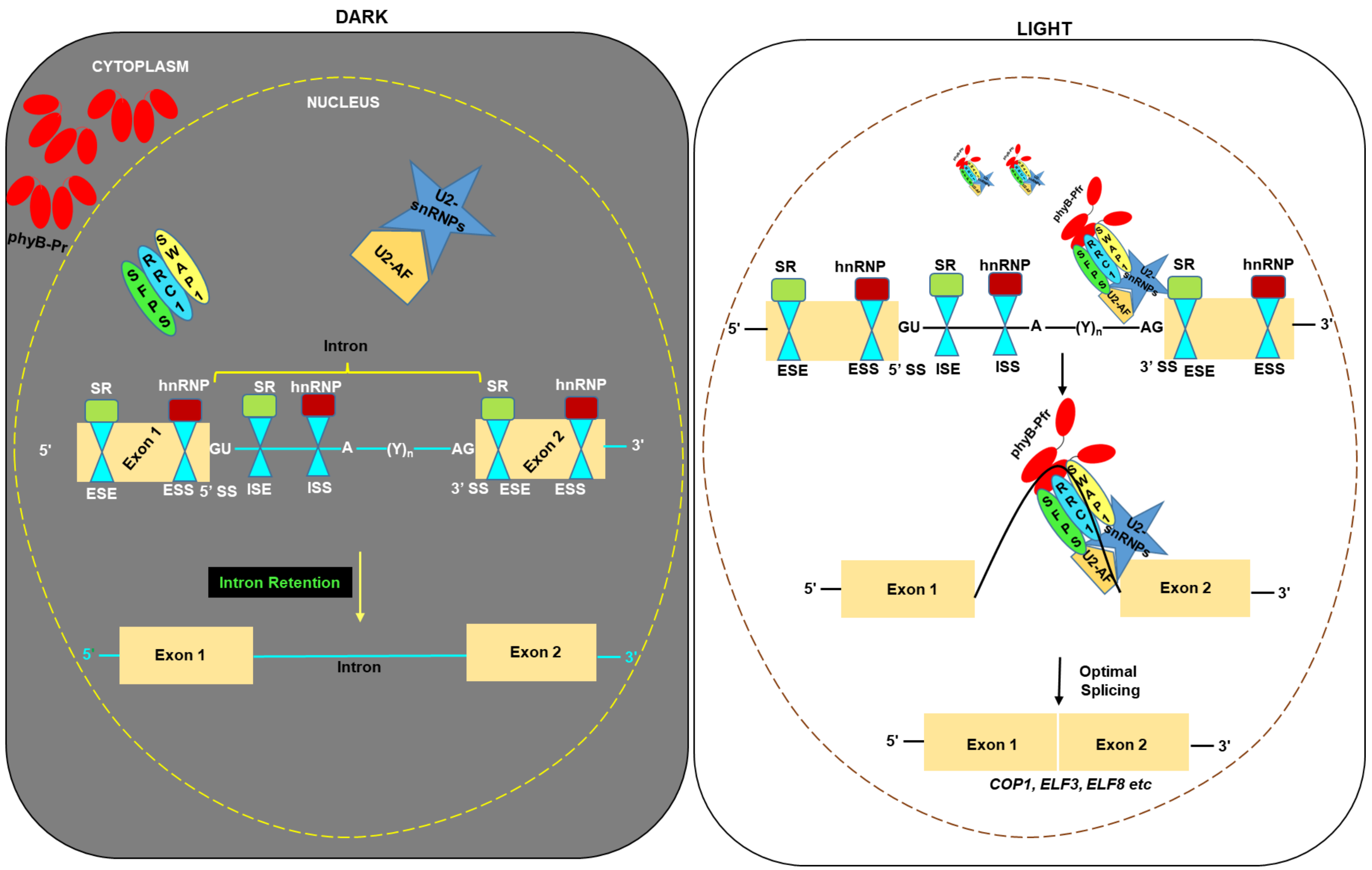


**Fig. S17: Model of red light modulated pre-mRNA splicing through SFPS-RRC1-SWAP1 ternary complex**

**Left Panel:** In the dark, biologically inactive phyB-Pr remains in the cytosol and therefore, does not interact with nuclei localized splicing factors or spliceosome complex. Nevertheless, splicing factors remain biologically active and target hundreds of pre-mRNAs to modulate their splicing in the dark. **Right Panel:** In response to the red-light irradiation, photoconverted and biologically active phyB-Pfr moieties migrate into the nucleus and interact with splicing factors. This possibly leads to certain biochemical changes within splicing factors, resulting in the targeting of a different set of pre-mRNAs to promote optimal photobiological responses in seedlings/plants.

Table S1: Primer sequences used in this study

| Primer | Sequence |
|---|---|
| SWAP1-Pro-F | CACC GGG TGA TTT TTT GCC ACC TCC T |
| SWAP1-Entry-F | CACC ATG GAT CGA AGA CAG CAC GA |
| SWAP1-R | TCT GGT TGT AGC TCT TGC ACT CA |
| SWAP1-Nhe1-F | AGA GCT AGC ATG GAT CGA AGA CAG CAC GAT TAT |
| SWAP1-FLAG1-R | ATC GTC TTT GTA GTC TCT GGT TGT A |
| SWAP1-FLAG2-R | GTC TTT GTA GTC CTT GTC GTC ATC GTC |
| SWAP1-FLAG3-R | TTT GTA GTC CTT GTC GTC ATC GTC TTT GTA |
| SWAP1-FLAG4-SmaI-R | ATA CCC GGG TCA CTT GTC GTC ATC GTC TTT GTA |
| SWAP1-BamH1-F | GCTGGATCCATGGATCGAAGACAGCACGA |
| SWAP1-Xba1-R | CCATCTAGATCTGGTTGTAGCTCTTGCACTC |
| SWAP1-EcoR1-F | AGA GAA TTC ATG GAT CGA AGA CAG CAC GAT TAT |
| SWAP1-Xma1-R | AGT CCC GGG C TCT GGT TGT AGC TCT TGC ACT CA |
| SWAP1-BamH1-R | GCTGGATCCTCATCTGGTTGTAGCTCTTGCA |
| SWAP1-qPCR-F | AAGTGGACCACCTGCAAATACC |
| SWAP1-qPCR-R | GTTGCAGCATTCCTGGACGTTG |
| ELF3-Intron1 3'-F | TTCTTGTGATTGCCCTGAGCAT |
| ELF3-Intron1 3'-R | TCCATGAAGGACATTTGGGAGACAA |
| ELF3-Intron2-F | CTGTCTCGGTTTGGTATTGCT |
| ELF3-Intron2-R | TTTAGATCCTGGGGTCCTCG |
| ELF3-Intron3-F | CAAACCTCTTCAACTGTGTAATAATCA |
| ELF3a-F1 | CCATTGCCAATCAACAAAGAG |
| ELF3a-R1 | TGGTCAGTCTTCTCCGAGTCAC |
| PP2A-Pro-F | TTA GCT GCT GCG AAA GAC GAG |
| PP2A-Pro-R | TTCCAAGTTCCGAGCGATCTATC |
| ELF3-Pro-F | AGC GAG TAT AAC CGT ATG ACC A |
| ELF3-Pro-R | GAGTGGTCGGATAGTGAGAAATC |
| BUD13-T-F | GGCAAAGGTTTAGCTCAGAAGCG |
| BUD13-T-R | TCAAGCTCCGGATCATCCCTTG |
| BUD13-IJ-F | ATT CAT AAA GAG ACC AGT CTT CA |
| BUD13-IJ-R | GAA TCC ATA TTT TTA AAC CTG TAG |
| RS31-T-F | AGGTCCAGCAGCTTATGAAAGACG |
| RS31-T-R | CTCTTGGGACTGGAGAACGACTTC |
| RS31-IJ-F | CTT CCT GCA AAA TCA TTT CTA CA |
| RS31-IJ-R | CGA TGA TTG GTA ATA GAC AGA C |
| BIM3-T-F | ATCTGTGCAGTTAAGCCTTCGG |
| BIM3-T-R | ACCATTATCCTCAGAAGCAAACGC |
| BIM3-IJ-F | TGG ATA CTA AGA GAG AAT GAT AT |
| BIM3-IJ-R | CAT TTA TCT CAT ACT CTA TCC TA |
| COP1-T-F | GTT GTA AAT GAA CCA GCA GAT AT |
| COP1-T-R | TTC ATG CTT ATT CCA ACT CAA GC |
| COP1-IJ-F | CAA TTA TTA CTA TGG GAT TCG TT |

| | |
|---|---|
| COP1-IJ-R | TCT CCT CAA GTG GCC TGC T |
| ELF3-T-F | CAA GTT TAT CTA AGT GTG GTT TA |
| ELF3-T-R | AGA AAC TTT CAT CGA ACT TCA GA |
| ELF3-IJ-F | GAG GGA CAT TCT CTG TTG ACC |
| ELF3-IJ-R | AAG CAA TAC CAA ACC GAG ACA G |
